# Elevated levels of iodide promote peroxidase-mediated protein iodination and inhibit protein chlorination

**DOI:** 10.1101/2024.02.23.581700

**Authors:** Kathrine V. Jokumsen, Valerie H. Huhle, Per M. Hägglund, Michael J. Davies, Luke F. Gamon

**Affiliations:** Dept. of Biomedical Sciences, University of Copenhagen, Copenhagen, Denmark; Department of Neuropathology, Charité – Universitätsmedizin Berlin, corporate member of Freie Universität Berlin and Humboldt Universität zu Berlin, Berlin, Germany

**Keywords:** Protein oxidation, post-translational modifications, myeloperoxidase, hypochlorous acid, chlorination, iodination, 3-iodotyrosine, 3-chlorotyrosine, methionine sulfoxide

## Abstract

At inflammatory sites, immune cells generate oxidants including H₂O₂. Myeloperoxidase (MPO), released by activated leukocytes employs H₂O₂ and halide/pseudohalides to form hypohalous acids that mediate pathogen killing. Hypochlorous acid (HOCl) is a major species formed. Excessive or misplaced HOCl formation damages host tissues with this linked to multiple inflammatory diseases. Previously (*Redox Biology*, 2020, 28, 101331) we reported that iodide (I⁻) modulates MPO-mediated protein damage by decreasing HOCl generation with concomitant hypoiodous acid (HOI) formation. HOI may however impact on protein structure, so in this study we examined whether and how HOI, from peroxidase/H₂O₂/I⁻ systems ± Cl⁻, modifies proteins. Experiments employed MPO and lactoperoxidase (LPO) and multiple proteins (serum albumins, anastellin), with both chemical (intact protein and peptide mass mapping, LC-MS) and structural (SDS-PAGE) changes assessed. LC-MS analyses revealed dose-dependent iodination of anastellin and albumins by LPO/H_2_O_2_ with increasing I⁻. Incubation of BSA with MPO/H_2_O_2_/Cl⁻ revealed modest chlorination (Tyr286, Tyr475, ∼4%) and Met modification. Lower levels of these species, and extensive iodination at specific Tyr and His residues (>20% modification with ≥10 µM I⁻) were detected with increasing I⁻. Anastellin dimerization was inhibited by increasing I⁻, but less marked changes were observed with albumins. These data confirm that I⁻ competes with Cl⁻ for MPO and is an efficient HOCl scavenger. These processes decrease protein chlorination and oxidation, but result in extensive iodination. This is consistent with published data on the presence of iodinated Tyr on neutrophil proteins. The biological implications of protein iodination relative to chlorination require further clarification.

## 1. Introduction

Iodide (I⁻) is an essential dietary mineral and plays a critical role in thyroid function and development [1]. This anion is concentrated in this gland via the actions of the Na^+^/I⁻ symporter, where it is used by a specialized heme peroxidase, thyroid peroxidase (TPO) together with hydrogen peroxide (H_2_O_2_) to generate reactive iodine species (variously reported as atomic iodine I.; iodinium ions, I^+^; or hypoiodous acid, HOI) that react with thyroglobulin at specific tyrosine (Tyr) residues [2, 3]. Subsequent processing of the iodinated species gives the thyroid hormones, thyroxine (T4, which contains 4 iodine atoms) and triiodothyronine (T3, the major metabolically active species) [2]. These hormones play a key role in regulating body mass, metabolism, temperature and organ growth (particularly the skin, hair and nails). Deficiencies in I⁻ in the diet, or bodily uptake, results in decreased levels of these hormones and enlargement of the thyroid gland (goitre), together with multiple clinical symptoms [2]. Auto-antibodies against TPO results in Hashimoto’s disease.

Whilst TPO shows a high specificity for oxidation of I⁻ [3], it can also oxidize other species to a limited extent, with this including the pseudohalide anion, thiocyanate (SCN⁻) [4]. This propensity for oxidation of multiple anions is shared by other members of the peroxidase superfamily, including lactoperoxidase (LPO), eosinophil peroxidase and myeloperoxidase (MPO) [5–7]. Each of these heme enzymes can oxidize I⁻ to reactive iodide-derived species such as HOI at appreciable rates [7], potentially resulting in protein modification by iodide-derived species, though all of these other peroxidases oxidize I⁻ to a lesser extent than other substrates [5]. Thus lactoperoxidase, generates high levels of HOSCN from oxidation of SCN⁻, eosinophil peroxidase oxidizes both Br⁻ (to give HOBr) and SCN⁻, and MPO oxidizes all of these species, together with chloride (Cl⁻) yielding hypochlorous acid (HOCl) [5–8]. The rates of oxidation of the different anions by these peroxidase enzymes various significantly, as do the concentrations of the anions in biological fluids [5–8]. As a consequence, the major oxidant(s) generated by each enzyme is situation dependent. In the case of MPO, the high concentration of Cl⁻ in most bodily fluids results in HOCl being a major oxidant, even though this anion is only slowly oxidized relative to the other anions (apparent second order rate constants for reaction of Compound I of MPO with these anions, at pH 7, are 2.5 × 10^4^, × 10^6^, 7.2 × 10^6^, and 9.6 × 10^6^ M^−1^ s^−1^ for Cl⁻, Br⁻, I⁻ and SCN⁻ respectively [7]). Evidence from studies using mixtures of anions (at physiologically-relevant concentrations) has indicated that plasma concentrations of SCN⁻, but not I⁻, can modulate the extent and nature of the damage generated by MPO, with a decrease in the extent of HOCl-mediated modifications [9]. The related selenium species selenocyanate (SeCN⁻) acts in a similar manner [10].

In contrast, *supplementation* studies using I⁻ indicate that elevated levels of this anion can minimize biological damage. This includes protection against oxidation of isolated proteins and protein mixtures [11], and amelioration of ischemia-reperfusion injury in mice, rats and pigs [12–14]. These effects may arise from modulation of HOCl formation and are based on the assumption that HOI (and HOSCN, when levels of SCN⁻ are increased by supplementation [15–18]) is less damaging than HOCl. This predication is supported by the E° values of these species, with HOCl having a higher redox potential (1.28 V) than HOI (0.78 V) [7]. Thus HOI might be expected to react less rapidly, and be more selective than HOCl. However, whether this translates into decreased biological effects is less obvious, as highly selective and specific modifications can be of major importance (cf. the role of specific oxidation reactions in cell signalling, and activation/deactivation of key biological pathways [19, 20]). In this context, the reactivity of HOI is relatively underexplored, especially when generated in competition with HOCl. In previous work, we observed dramatic changes in protein oxidation caused by MPO at low physiologically relevant concentrations of I⁻ (< 5 µM) when used in combination with physiological Cl⁻ concentrations (100 mM) [21]. Even though I⁻ is preferred as a substrate by MPO over Cl⁻ (with a specificity constant 288 fold higher than Cl⁻ [22]), this effect occurs at concentrations much lower than would be predicted based on reaction rates alone [21]. Furthermore, a recent study has provided evidence for the presence of iodinated proteins in activated human neutrophils, in sputum from people with cystic fibrosis and from skin abscesses from people infected with *S. aureus* [23]. These iodinated species were more effective than their chlorinated analogues in activating chemotaxis and inducing cytokine release, providing strong evidence for the occurrence of both widespread protein iodination in biological samples, and potent effects of these species [23].

In the light of these data, this work examined the reactivity of enzyme-generated HOI, both when generated in isolation (using a LPO/H₂O₂ system) and in competition with HOCl by a MPO/H₂O₂/l⁻/Cl⁻ system. We demonstrate that HOI preferentially targets tyrosine (Tyr) and to a lesser extent histidine (His) residues on proteins (forming iodo-tyrosines and iodo-histidines) and that these products dominate over HOCl-derived species (3-chlorotyrosine and methionine sulfoxide). Furthermore, these reactions occur with very low concentrations of I⁻ added to MPO/H₂O₂/Cl⁻ systems. This study is the first comprehensive study of HOI-derived protein modification and provide important data on the potential formation of iodinated products by MPO systems at sites of inflammation.

## 2. Materials and methods

### 2.1. Chemicals and reagents

All chemicals were of high purity and purchased from Sigma-Aldrich (Søborg, Denmark) unless otherwise stated. All solutions for SDS-PAGE and LC-MS were prepared with nano-pure H_2_O sourced from a Millipore MilliQ system. Cl⁻ and I⁻ solutions were prepared using NaCl and KI in MilliQ H_2_O. Phosphate-buffered saline (PBS) 20X was purchased from VWR (Søborg, Denmark). SDS-PAGE was carried out using NuPAGE 4-12% Bis-Tris gel 1.0 mm 12 well (NP0322BOX), NuPAGE MES SDS Running Buffer 20X (NP0002), NuPAGE LDS Sample Buffer 4X (NP0007), and NuPAGE Sample Reducing Agent 10X (NP0009) which were purchased from Thermo Fisher (Roskilde, Denmark). Precision Plus Protein Kaleidoscope Standards (1610375) were obtained from BioRad (Copenhagen, Denmark). Recombinant anastellin was a kind gift from Assoc. Prof. Pontus Gourdon (Dept. Biomedical Sciences, Univ. of Copenhagen) and produced as described previously [24]. Sequencing grade modified trypsin (V5111), rLys-C (V1671), and Glu-C (V1651) were obtained from Promega (Finnboda, Sweden).

### 2.2. Oxidation of proteins for subsequent SDS-PAGE analysis

Human, bovine, or mouse serum albumins (3, 3 and 5 µM final concentration, respectively) were treated with either an LPO/H₂O₂/I⁻ or MPO/H_2_O_2_/Cl⁻/I⁻ system in 100 mM chelex-treated phosphate buffer, pH 5.8 for LPO and pH 7.4 for MPO. Anastellin (5 µM final concentration) was treated with an MPO/H_2_O_2_/Cl⁻/I⁻ system in chelex-treated phosphate buffer, pH 7.4. For human serum albumin treated with LPO, the H₂O₂ concentration was varied between 3 - 1500 µM while the I⁻ concentration was kept constant at 10 mM, or the I⁻ concentration was varied in a range of 0.01 - 10 mM while the H₂O₂ concentration was constant at 3 µM. H₂O₂ was added in five aliquots over a period of 10 min to samples containing 1.5 µM LPO and I⁻ followed by incubation for 2 h at 37°C. For mouse serum albumin, bovine serum albumin or anastellin treated with LPO or MPO, H_2_O_2_ concentration was kept constant at 50 or 75 µM while the I⁻ concentration was varied in a range of 0.1 - 1000 µM. H_2_O_2_ was added as a bolus addition to samples containing 1.5 µM LPO or 100 nM MPO, followed by incubation for 2 h at 37°C.

Samples were prepared for SDS-PAGE using NuPAGE LDS Sample Buffer in a 1:4 dilution to the final volume. NuPAGE Sample Reducing Agent was added in a 1:10 dilution to the reduced samples, while H_2_O was added in an equal volume to the non-reduced samples. The protein samples were heat denatured for 10 min at 70 °C, and 1 µg of protein was then loaded per well. Precision Plus Protein Kaleidoscope Standards were included as a reference for protein masses. SDS-PAGE was carried out using a NuPAGE 4-12% Bis-Tris gels which was run at 150 V for 1 h in NuPAGE MES SDS Running Buffer. Afterwards, the proteins on the gel were visualized by silver staining. Thus, the gel was fixed in 50% methanol / 10% acetic acid in H_2_O for 30 min, rinsed in 5% methanol in H_2_O for 15 min, rinsed 3x in H_2_O for 5 min, sensitized in sodium thiosulfate in H_2_O (0.02% w/v) for 2 min, rinsed in H_2_O for 2 min, stained in silver nitrate in H_2_O (0.2% w/v) for 25 min, and rinsed in H_2_O for 5 min. The stain was developed for 1 - 2 min in sodium carbonate (3% w/v) / formaldehyde (0.0185% v/v) / sodium thiosulfate (0.0004% w/v) in H_2_O until the stop solution of EDTA (1.4% w/v) in H_2_O was added followed by a wash in H_2_O for 5 min.

### 2.3. LC-MS based intact protein analysis of anastellin modified by a lactoperoxidase (LPO) system

Anastellin (5 µM final concentration) was incubated with an LPO/H₂O₂/I⁻ system in 100 mM chelex-treated phosphate buffer, pH 5.8. Five aliquots of H₂O₂ (5 x 10 µM; 50 µM total) were added over a period of 10 min to samples containing LPO (1.5 µM) and I⁻ (1 - 1000 µM) prior to incubation at 37 °C for 2 h. The samples were purified by solid-phase extraction using C4 Stagetips (2 layers of Empore^TM^ SPE disks C4) as previously described[25]. The Stagetips were conditioned with 100% methanol and equilibrated with 0.1% trifluoroacetic acid (TFA) in H_2_O. The samples were diluted and acidified with 1% TFA in H_2_O (100 µL final volume, 0.5% TFA final concentration) to improve ion-pairing and binding of the protein to the C4 material. Samples were loaded into the tips and washed with 0.1% TFA in H_2_O followed by elution with 50% and 80% acetonitrile (ACN) containing 0.1% TFA. The elutes were dried at 60 °C under vacuum for 1 h and then resuspended in 20 µL 0.1% formic acid (FA) in H_2_O.

Intact anastellin was analysed by using a Dionex Ultimate 3000 liquid chromatography (LC) system (Thermo Fisher Scientific) coupled online to an Impact HD II mass spectrometer (Bruker, Bremen, Germany) with electrospray ionization, in positive ion mode. 5 pmol protein were injected and separated by gradient elution at a flow rate of 10 µL min^-1^ using a Thermo Scientific MAbPac^TM^ 4 µm Reversed Phase 150 x 0.15 mm HPLC column. Elution was started at 5% buffer B for 3 min, followed by a gradient elution from 5% to 60% B over 12 min and 60-95% B over 3 min. Buffer B was kept at 95% for 9 min before it was decreased to 5% B over 1 min and re-equilibrated with 5% B for 2 min. Buffer A consisted of 0.1% FA in water and buffer B contained 0.1% FA and 80% ACN in water. The data were analysed manually using DataAnalysis and QuantAnalysis (Bruker) with different intact species quantified from the most abundant charge state (+13), relative to the unmodified anastellin peak.

### 2.4. Oxidation and digestion of anastellin for subsequent LC-MS based peptide mass mapping

Anastellin (5 µM) incubated for 2 h at 37 °C with MPO (100 nM), H_2_O_2_ (50 µM), NaCl (100 mM), KI (0, 10 µM, 100 µM or 1 mM) in 100 mM chelex-treated phosphate buffer, pH 7.4 in a reaction volume of 20 µL, was subjected to the Solid-Phase Protein Preparation (SP^3^) protocol [26]. Briefly to samples were added 40 µL MilliQ H_2_O, 2 µL of a 1:1 mix of two types of commercially available carboxylate beads (Sera-Mag Speed beads, 45152105050250, 65152105050250, GE Healthcare), and 60 µL ethanol. The bead/protein mixture was incubated in a thermal mixer at 24 °C at 1000 rpm for 30 min before the tubes were placed in a magnetic rack and the supernatants removed. The bead/protein mixture was washed twice with 80 % ethanol in H_2_O, then 100 µL of 50 mM Tris-HCl (pH 7.5) and 1 µL trypsin (100 ng µL^-1^) were added, and incubated overnight at 21 °C. The next day, 1 µL Glu-C (0.1 µg µL^-1^) was added, samples were incubated for 4 h at 21 °C, and the supernatants removed after placing the tubes in the magnetic rack, and subsequently subjected to solid-phase extraction using C18 Stagetips [25]. Samples were diluted and acidified with 10% TFA in H_2_O (1% TFA final concentration) to improve binding of the peptides to the C18 material and then loaded onto the tips. The Stagetips were washed with 0.1% TFA in H_2_O followed by elution with 80% ACN in H_2_O containing 0.1% TFA. The eluates were dried under vacuum and then reconstituted in 20 µL 0.1% FA in H_2_O. The samples were subjected to ESI-MS as described in Section 2.3., but with 5 µL (25 pmol) of anastellin digest loaded on a 150 x 0.5 mm Kinetex 2.6 µm XB-C18 column (Phenomenex) operating at 40 °C with a flow rate of 30 µL min^-1^. Peptides were eluted over a 20 min linear gradient from 2.5 to 45% buffer B (0.1% FA, 80% ACN in H_2_O), with 0.1% FA in H_2_O as buffer A.

### 2.5. Oxidation and digestion of serum albumins for subsequent LC-MS based peptide mass mapping

Mouse or bovine serum albumin (5 and 2 µM final concentration, respectively) was treated with either an LPO/H₂O₂/I⁻ or MPO/H_2_O_2_/Cl⁻/I⁻ system in 100 mM chelex-treated phosphate buffer, pH 5.8 for LPO, and pH 7.4 for MPO. The LPO/H₂O₂/I⁻ system consisted of 1.5 µM LPO, 50 µM H₂O₂ (10x molar excess relative to protein, bolus addition), and I⁻ over a concentration range of 1 - 500 µM. The MPO/H₂O₂/I⁻/Cl⁻ system consisted of 100 nM MPO, 20 µM H₂O₂ (10x molar excess relative to protein, bolus addition), 100 mM Cl⁻, and I⁻ in a concentration range of 1 - 500 µM. The proteins were incubated with the oxidant systems for 2 h at 37 °C, followed by reduction and alkylation with tris-(2-carboxyethyl) phosphine (TCEP) and chloroacetamide (CAA) (1:10 of final volume) under denaturing conditions in 8 M urea / 100 mM Tris (pH 8) for 30 min at 21 °C. Digestion of the protein was carried out on magnetic beads as described in Section 2.4.. The beads (15:1, beads:protein) were added to the samples along with ACN (70% final concentration v/v in H_2_O) and incubated for 20 min at 21 °C. Samples were then placed on a magnetic rack, supernatants were discarded, and the beads were washed twice with 70% ethanol in H_2_O and once with 100% ACN. The samples were left to dry for 10 min in order for ACN to evaporate before the digestion buffer (100 mM Tris, pH 8) was added containing protease enzymes. Both proteins were digested with trypsin (1:50, enzyme:substrate) overnight at 37 °C. In separate experiments, mouse serum albumin was subjected to a two-step digestion where the protein first was incubated with Lys-C (1:20, enzyme:substrate) for 1 h at 37 °C followed by the addition of a digestion buffer (100 mM triethylammonium bicarbonate) containing Glu-C (1:20, enzyme:substrate) and incubation overnight at 37 °C. Regardless of the enzyme digestion performed, the samples were subsequently placed on magnetic racks and the supernatants containing the peptides were collected. An additional elution was carried out using 2% dimethyl sulfoxide (DMSO) in H_2_O, and the two supernatants were pooled together. The samples were purified by solid-phase extraction using C18 Stagetips (2 layers of Empore^TM^ SPE disks C18) that were conditioned and equilibrated as described above. The Stagetips were washed with 0.1% TFA in H_2_O followed by elution with 50% ACN containing 0.1% TFA in H_2_O. The eluates were dried at 60 °C under vacuum for 1 h and then resuspended in 25 µL 0.1% formic acid (FA) in H_2_O.

### 2.6. LC-MS based peptide mass mapping analysis of serum albumins

MSA and BSA digests were analysed by ESI LC-MS in the same general manner as described above for intact protein analysis (Section 2.3.), with minor changes. Four pmol protein digest was injected into the instrument and separated by gradient elution using either a Phenomenex Luna® LC column (MSA; 5 µm C18(2) 250 x 4.6 mm) or a reversed-phase C18 column with an integrated CaptiveSpray Emitter (BSA; 25 cm × 75 µm; 1.6 µm particle size; IonOpticks) for reversed-phase chromatography using a Dionex Ultimate 3000 system (Thermo Fisher Scientific). For the MSA digests, peptides were separated at a flow rate of 20 µL min^-1^ using buffer A (0.1% FA in H_2_O) with 5 to 40% buffer B (0.1% FA, 80% ACN in H_2_O) over a 27 min linear gradient elution, followed by 40-95% B over 2 min. Buffer B was kept at 95% for 5 min before it was decreased to 5% B over 1 min and re-equilibrated with 5% B for 3 min. For the BSA digests, peptides were separated at a flow rate of 400 nL min^−1^ over a period of 100 min with a 3-step binary gradient with mobile phases A and B comprising 0.1% v/v FA in H_2_O, and 0.1% v/v FA in ACN, respectively. Step 1 consisted of an increasing gradient of B of 2% to 15% over 60 min, step 2 was an increasing gradient of B from 15% to 25% over 30 min, and step 3 an increasing gradient of B from 25% to 37% over 10 min. The column was then washed with 20% A and 80% B, followed by re-equilibration with 98% A and 2% B.

### 2.7. Relative quantification of oxidative modifications by mass spectrometry

Mass spectrometry data were analysed using Maxquant [27], Fragpipe [28] and Skyline [29]. Database searches were performed using the protein sequence of MSA as reference (UniProt no. Q546G4) or the recombinant sequence of anastellin (Supplementary Figure 7) to identify peptides based on their MS and MS/MS spectra. In the search settings variable modifications were set to Met and Trp oxidation (+16 Da), Trp dioxidation (+32 Da), Tyr, Trp and His chlorination (+33.96 Da) and dichlorination (67.92 Da), as well as Tyr, Trp and His iodination (+125.9 Da) and Tyr diiodination (+251.8 Da). Adding these modifications to the database search means that all MS/MS spectra that were not identified as MSA digest peptides are submitted to another search level where a hypothetical modification is positioned on each amino acid and a score is given based on the mass difference between the identified and unidentified MS/MS spectra [27]. Database searches using MaxQuant (version 1.6.1.0) were performed with tryptic constraints, one missed cleavages, 1% peptide and protein level false discovery rates, carbamidomethylation of Cys as a fixed modification and mass accuracy of 20 ppm. Database searches using Fragpipe (version 16.0, MSfragger version 3.3, Philosopher version 4.1.1 [30]) were performed with tryptic or Lys-C/Glu-C constraints (for MSA) or trypsin/GluC (for anastellin), two missed cleavages, 1% peptide and protein level false discovery rates, carbamidomethylation of Cys as a fixed modification and mass accuracy of 20 ppm.

The search data were then transferred to Skyline, where all identified peptides were quantified based on the extracted ion chromatogram of two precursor ions (e.g. precursors for the +2 and +3 charge state). All identifications were validated by manual inspection of the chromatogram considering the peak shape, retention time, mass error, and isotopic distribution, and, in some cases, checking the quality of the MS/MS spectra. The extent of modification (% occupancy) at each Tyr residue was calculated by dividing the signal intensity (area-under-the-curve) of the peptide containing the modified Tyr with the sum of the signal intensity of all identified forms (non-modified and modified) of this peptide with the value expressed as a percentage. If more than one peptide contained a particular Tyr, the percentage modification was determined as the mean of the percentage modification calculated for each individual peptide as described above. If a peptide contained multiple modified amino acids, the percentage modification on each residue was calculated as the sum of signal intensities of all forms of the peptide containing the modification divided by the summed signal intensity of all identified peptide forms. A few peptides contained two Tyr residues that could be modified. In some cases, two peaks were visible in the extracted ion chromatogram of the modified peptide, with this ascribed to the presence of two different species generated by modification of either of the two Tyr residues. These peaks were quantified separately and randomly assigned to one of the Tyr residues, since it was not possible to determine which peak belonged to which Tyr from the Maxquant search or MS/MS spectra. In other cases, the peaks from the two different isomers were not separated in the chromatogram and the percentage modification was split equally between the two Tyr residues.

The loss of parent ion was quantified by taking the peak area ratio, which was calculated by dividing the signal intensity of the unmodified peptide with the sum of the signal intensity of all peptides in the sample, and dividing it by the peak area ratio of the 0 µM I⁻ control. In cases where the peptide contained an oxidized Met, the peak area ratio was calculated as the sum of the signal intensity of the unmodified and Met oxidized peptide species, divided by the sum of the signal intensity of all peptides in the sample to evaluate the effect of Tyr modification alone.

### 2.8. Statistical analysis

All analyses were performed using R Statistical Software (v4.3.2; R Core Team 2023) with *p* < 0.05 considered significant. One-way analysis of variance (ANOVA) with post hoc Dunnett’s test was used to compare the effects of increasing I⁻ concentrations with a control (no I⁻). Two-sample t-test was used to compare MPO/H_2_O_2_/Cl⁻ with no I⁻, to an untreated control.

## 3. Results

### 3.1. The enzymatic activity of lactoperoxidase induces multiple iodinations on anastellin as detected by intact protein LC-MS analysis

As the chemistry of HOI is relatively underexplored, we assessed the reactivity of this species, in the absence of other hypohalous acids, with a series of model proteins including MSA, BSA and anastellin. Anastellin is derived from the major extracellular matrix protein fibronectin [31], which is an established peroxidase target [32]. A LPO/H₂O₂/I⁻ system was employed to produce HOI, as this enzyme does not oxidize Cl^-^ [33]. Anastellin or MSA were treated with LPO/H₂O₂ and increasing I^-^ (0 - 1000 µM). LC-MS analyses of modified anastellin samples yielded spectra with ions with characteristic *m/z* changes of +125.9 Da assigned to loss of one H atom and addition of one I atom (Fig. 1A). These ions were detected even with very low levels of added I⁻ in the reaction system, consistent with highly efficient iodination (Fig. 1B). With increasing I⁻ concentrations, anastellin-derived species were detected with up to 20 iodine atoms per protein molecule, where species with up to 12 iodine atoms could be quantified (Fig. 1C).

**Figure 1.**
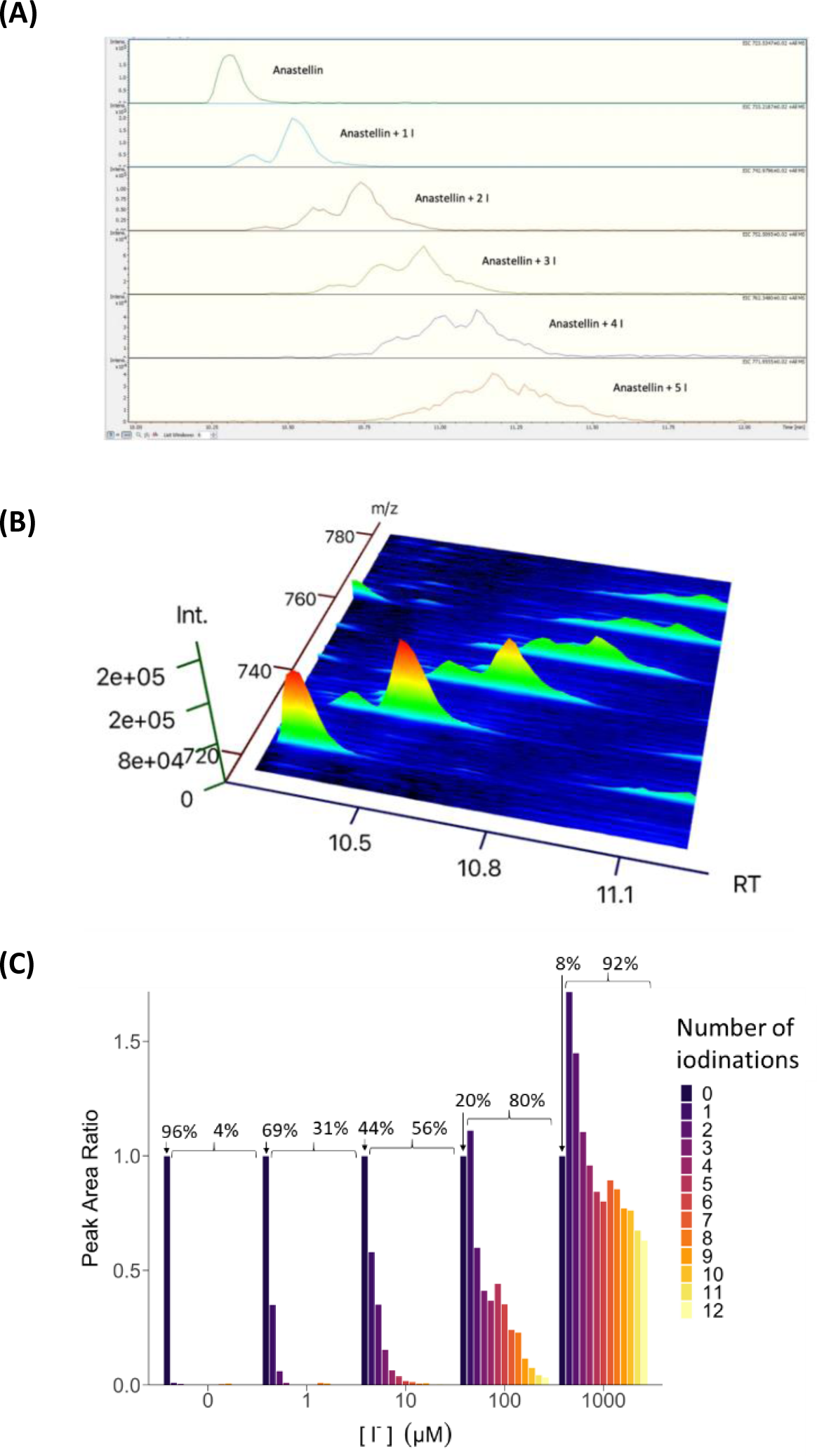
LC-MS intact protein analysis of unmodified and iodinated species of anastellin following treatment with LPO/H_2_O_2_/I⁻ system. Anastellin (5 µM) was treated with LPO (1.5 µM) and H_2_O_2_ (50 µM, 10-fold molar excess over anastellin concentration) in the absence or presence of increasing I⁻ (1 - 1000 µM) for 2 h at 37 °C in chelexed phosphate buffer, pH 5.8. Intact protein mass spectra (A) and (B) revealed iodinated species with a shift in mass and retention time. (C) The extent of iodination (for 1 - 12 added iodine atoms) was quantified relative to unmodified anastellin. Representative data from a single experiment of three replicates. Values above the bars in panel C, represent the % values for the indicated species of the total peak area.

Iodination resulted in a systematic increase in the retention time of the modified species on the chromatographic column (Fig. 1B; ∼0.2 min per additional iodine atom) with this assigned to an increased hydrophobicity of the iodine adducts. The formation of these iodinated species was not observed in the absence of LPO, or absence of H₂O₂. Modest levels of non-enzymatic iodination (from direct reaction of H_2_O_2_ with I⁻) were only observed with 1 mM I⁻ and 50 µM H_2_O_2_, which far exceed physiological levels [34]. Thus species generated enzymatically by LPO appear to be responsible for the facile protein iodination. Ions consistent di-oxidation (+31.99 Da; +2O) were also observed, though these were minor products. Similar studies with albumins exposed to the same LPO system revealed a large distribution of modified species with overlapping envelopes of peaks (Supplementary Fig. 1) but a lack of resolving power of the mass spectrometer, due to the higher molecular mass of these proteins, precluded detailed analysis and quantification.

### 3.2. The enzymatic activity of lactoperoxidase induces multiple iodinations and oxidations on serum albumins as detected by peptide mass mapping

To obtain data on the potential sites of modification on mouse serum albumin (MSA) exposed to the LPO system, these proteins were incubated with LPO/H₂O₂/I⁻ and subsequently digested with trypsin and Glu-C/Lys-C to release native and modified peptides for LC-MS (proteomic) analysis. This approach allows determination of the precise sites of modification, and quantification of the extent of both product formation and parent amino acid loss.

Initial experiments employed MSA (5 µM) treated with LPO (1.5 µM), H₂O₂ (10x molar excess, 50 µM total, bolus addition) and increasing concentrations of I⁻ (1 - 500 µM). After incubation, the samples were subject to tryptic digestion and analysis by LC-MS/MS. This resulted in a combined sequence coverage of 94.2%. All Tyr, Trp, and Met residues in the protein sequence (UniProt Q546G4) were covered (Supplementary Figure 2). Peptide MS/MS spectra were searched for potential ions arising from Tyr and Trp mono- (*m/z* +125.9 Da; M – H + I) and di-iodination (*m/z* +251.8 Da; M −2H + 2I), as well as incorporation of one or more oxygen atoms into Met and Trp (*m/z* +16 for Met, Trp; or + 32 Da for Trp). These searches resulted in the detection of species with multiple different types of modifications, including ions arising from Tyr iodination (Fig. 2A). Four major types of modifications were detected: oxidation at Met residues to species assigned to the sulfoxide (at 6 out of 7 residues in the total sequence; Table 1), mono- and di-oxidation at the single Trp in the protein (W238), and mono- and di-iodination of Tyr residues. Mono-iodination was detected at 16 of the 22 Tyr residues in the sequence (Table 1) whereas di-iodination was detected at 15 sites (Table 1). These ions are assigned to the presence of 3-iodotyrosine (3-ITyr) and 3,5-di-iodotyrosine (3,5-di-ITyr) respectively. The extent of modification at each site varied significantly (Fig. 2B,C), indicating markedly different susceptibilities to oxidation (i.e. a high degree of site-specificity), and a strong dependence on the I⁻ concentration.

**Figure 2.**
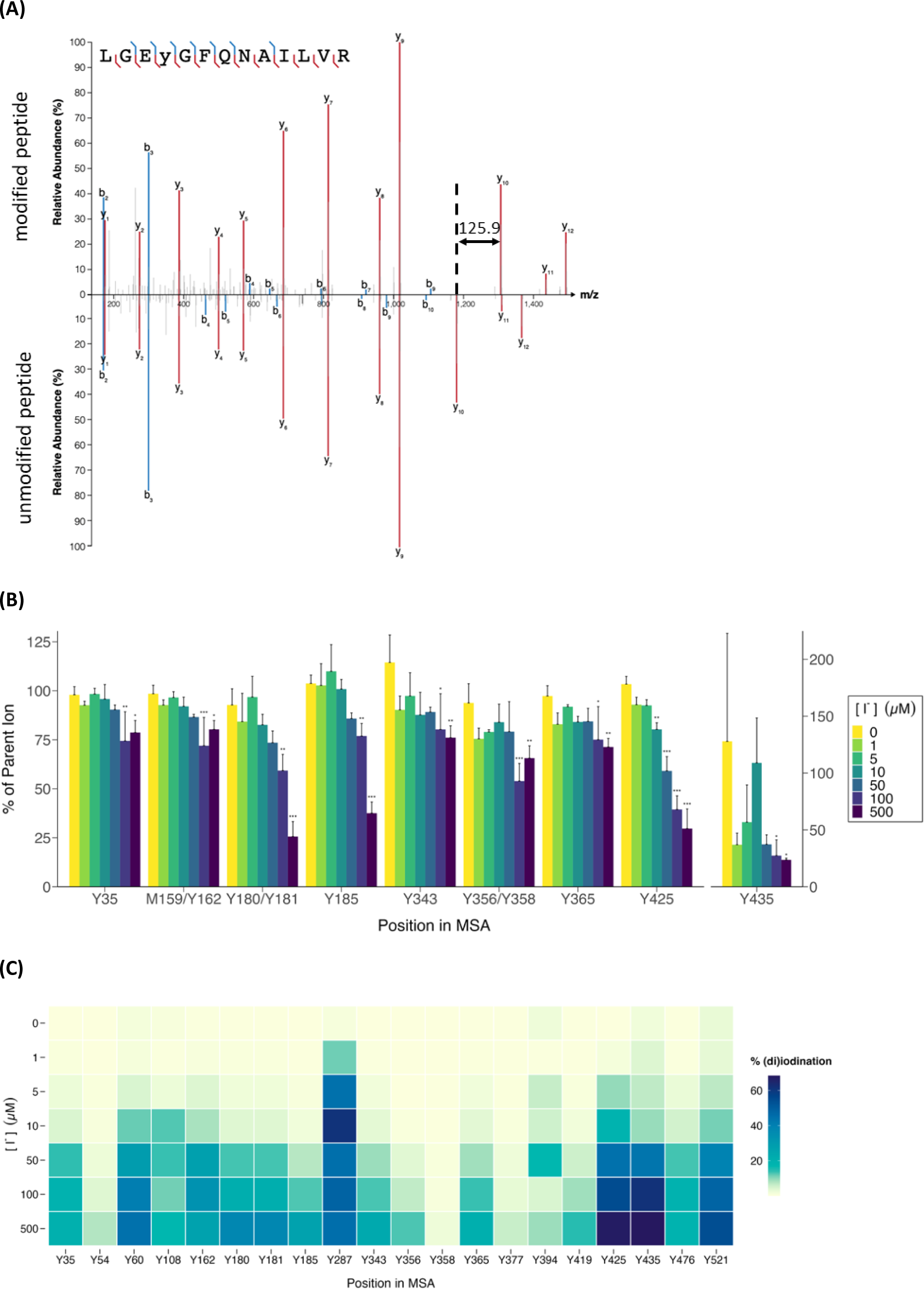
Mouse serum albumin (5 µM) was treated with LPO (1.5 µM) and H_2_O_2_ (10x excess, 50 µM) in the absence or presence of increasing I⁻ concentrations (1 - 500 µM) for 2 h at 37 °C before reduction/alkylation, digestion and LC-MS/MS peptide mapping. (A) MS/MS spectra from modified (top spectrum) and native (bottom spectrum) of the peptide _422_LGEYGFQNAILVR_434_ from MSA. Iodination of Y425 results in +125.9 Da mass shift of the y_10_ ion. (B) Sensitivity of different Tyr (Y) residues in MSA to modification by LPO/H_2_O_2_/I⁻ system, with only residues that were detected as significantly modified relative to the parent ion being presented as % values. The peptides containing M159/Y162, Y180/181 and Y356/358 have multiple potential reaction sites, and modification of both residues in each pair are expected to contribute to parent ion loss. (C) Heat map showing percentage iodination/di-iodination of all Tyr residues, except Y172 and Y174 which were not identified as being modified. Data are presented as mean + SD from 3 independent experiments Statistical significance was determined by one-way ANOVA with a post-hoc Dunnett’s test. * *p* < 0.05, ** *p* < 0.01, *** *p* < 0.001.

**Table 1.**
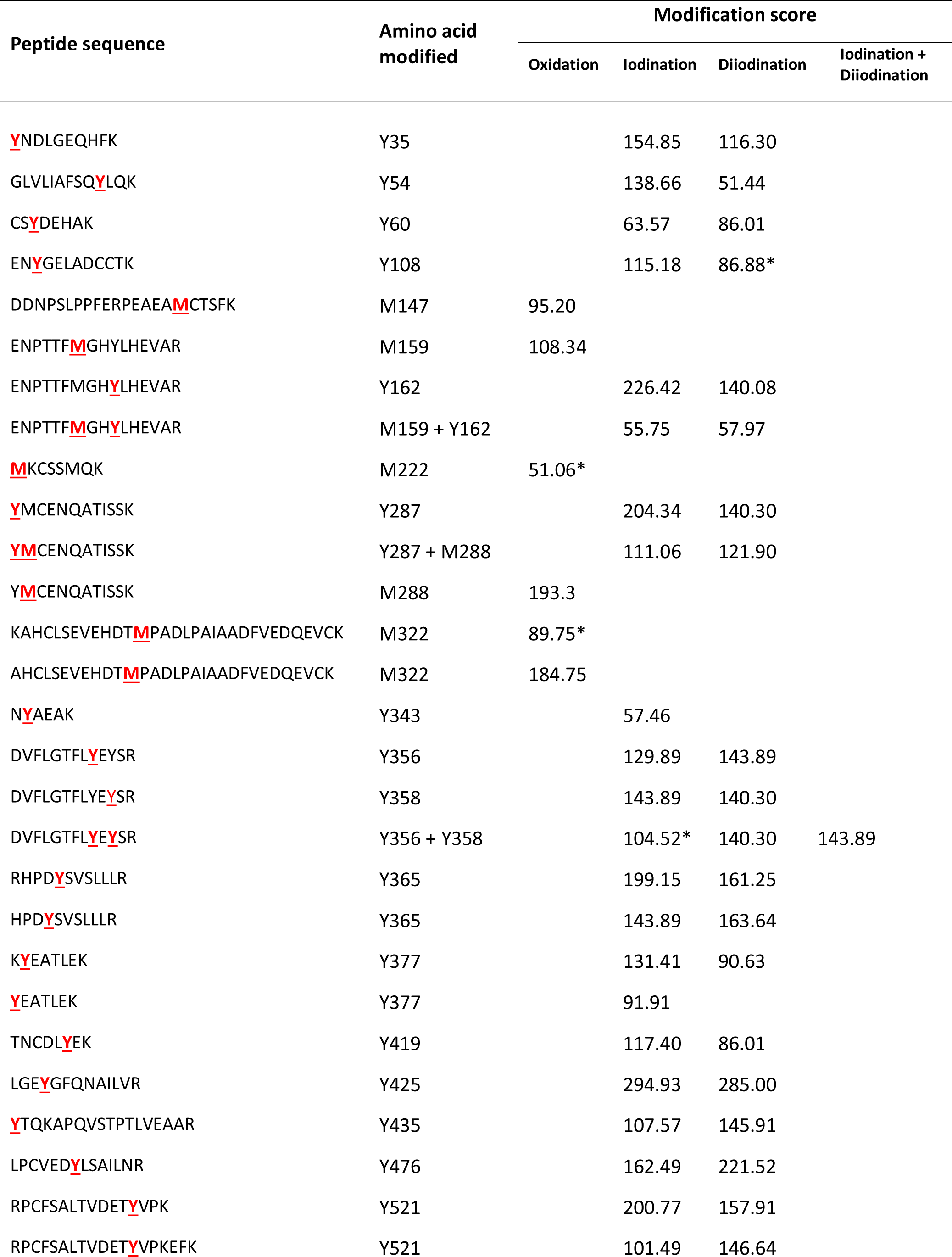
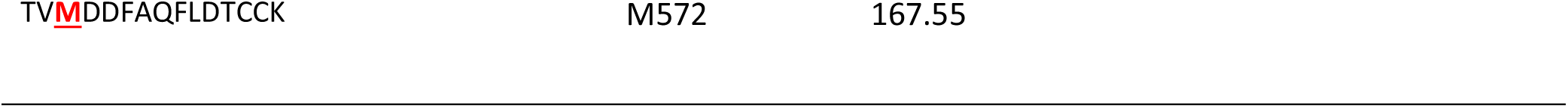
Nature and sites of Met oxidation and Tyr iodination and di-iodination induced by an LPO/H₂O₂/I⁻ system on mouse serum albumin (MSA) detected by LC-MS following reduction, alkylation, trypsin digestion and peptide mass mapping. Modified amino acid(s) are indicated in red and underlined in the peptide sequence. Numbering is from the protein sequence predicted from the gene with amino acids 1-18 corresponding to the signal peptide and amino acids 19-24 being the propeptide. ‘Modification score’ is a match score assigned to each modified peptide identified by Maxquant. All MS/MS spectra that were not identified as unmodified peptides generated by trypsin digestion of MSA, were submitted to another search level where hypothetical modifications were positioned on each amino acid. A match score was then given based on the mass difference between the identified and unidentified MS/MS spectra. * indicates that the peptide was not validated, based on the peak shape, retention time, mass error, isotopic distribution, and MS/MS spectra, and quantified in Skyline.

| Peptide sequence | Amino acid modified | Modification score |  |  |  |
| --- | --- | --- | --- | --- | --- |
|  |  | Oxidation | Iodination | Diiodination | Iodination + Diiodination |
| **Y**NDLGEQHF~~K~~ | Y35 |  | 154.85 | 116.30 |  |
| GLVLIAFSQ**Y**LQ~~K~~ | Y54 |  | 138.66 | 51.44 |  |
| CS**Y**DEHAK | Y60 |  | 63.57 | 86.01 |  |
| EN**Y**GELADCCTK | Y108 |  | 115.18 | 86.88* |  |
| DDNPSLPPFERPEAE**M**CTSF~~K~~ | M147 | 95.20 |  |  |  |
| ENPTTF**M**GHYLHEVAR | M159 | 108.34 |  |  |  |
| ENPTTFM**GHY**LHEVAR | Y162 |  | 226.42 | 140.08 |  |
| ENPTTF**M**GH**Y**LHEVAR | M159 + Y162 |  | 55.75 | 57.97 |  |
| **M**KCSSMQK | M222 | 51.06* |  |  |  |
| **Y**MCENQATISSK | Y287 |  | 204.34 | 140.30 |  |
| **YM**CENQATISSK | Y287 + M288 |  | 111.06 | 121.90 |  |
| **YM**CENQATISSK | M288 | 193.3 |  |  |  |
| KAHCLSEVEHDT**M**PADLPAIAADFVEDQE~~VCK~~ | M322 | 89.75* |  |  |  |
| AHCLSEVEHDT**M**PADLPAIAADFVEDQE~~VCK~~ | M322 | 184.75 |  |  |  |
| N**Y**AEAK | Y343 |  | 57.46 |  |  |
| DVFLGTFL**Y**EYSR | Y356 |  | 129.89 | 143.89 |  |
| DVFLGTFL**Y**EYSR | Y358 |  | 143.89 | 140.30 |  |
| DVFLGTFL**Y**E**Y**SR | Y356 + Y358 |  | 104.52* | 140.30 | 143.89 |
| RHPD**Y**SVSLLLR | Y365 |  | 199.15 | 161.25 |  |
| HPD**Y**SVSLLLR | Y365 |  | 143.89 | 163.64 |  |
| K**Y**EATLEK | Y377 |  | 131.41 | 90.63 |  |
| **Y**EATLEK | Y377 |  | 91.91 |  |  |
| TNCDL**Y**EK | Y419 |  | 117.40 | 86.01 |  |
| LGE**Y**GFQNAILVR | Y425 |  | 294.93 | 285.00 |  |
| **Y**TQKAPQVSTPTLVEAAR | Y435 |  | 107.57 | 145.91 |  |
| LPCVED**Y**LSAILNR | Y476 |  | 162.49 | 221.52 |  |
| RPCFSALTVEDET**Y**VPK | Y521 |  | 200.77 | 157.91 |  |
| RPCFSALTVEDET**Y**VPKEFK | Y521 |  | 101.49 | 146.64 |  |

The Tyr residues present at positions 287 and 425 (Y287, Y425) were highly sensitive to iodination and were modified to a significant extent even with low concentrations of added I^-^. Thus Y287 was detected as modified (% occupancy) to 11.4 ± 0.2 %, when 1 µM I⁻ was employed, and Y425 was modified to 9.3 ± 0.8 % with 5 µM I⁻. In contrast, other Tyr residues, such as Y54 and Y358, were not significantly iodinated until high I⁻ concentration were employed (Y54: 2.2 ± 0.1 % at 50 µM I⁻, Y358: 1.7 ± 0.4 % at 500 µM I⁻). Mono-iodination predominated over di-iodination at low I⁻ concentrations, but the extent of di-iodination increased markedly, for some residues, with high I⁻. These data are consistent with sequential mono- and di-modification. The detection of significant levels of di-iodination at some sites, whilst other Tyr remained essentially unmodified, suggests that the rate of addition of a second iodine atom to a Tyr which is already mono-iodinated is facile. The amount of iodination on each Tyr residue increased consistently, but not in a linear manner, with I⁻ concentration, reaching combined mono-/di-iodination yields as high as 70%.

Complementary analyses of the parent Tyr-containing peptides confirmed that most of the Tyr residues in the protein are significant targets of modification, although no significant loss was detected for Y54 and Y358 (Supplementary Figure 3), consistent with the low levels of iodination observed at these two residues (Fig. 3C). These data also confirmed the high extents of modification detected for Y287 and Y425, with ∼10% loss of the parent peptides detected with I⁻ concentrations as low as 1 µM. This value is within the known plasma I⁻ physiological range [34, 35].

**Figure 3.**
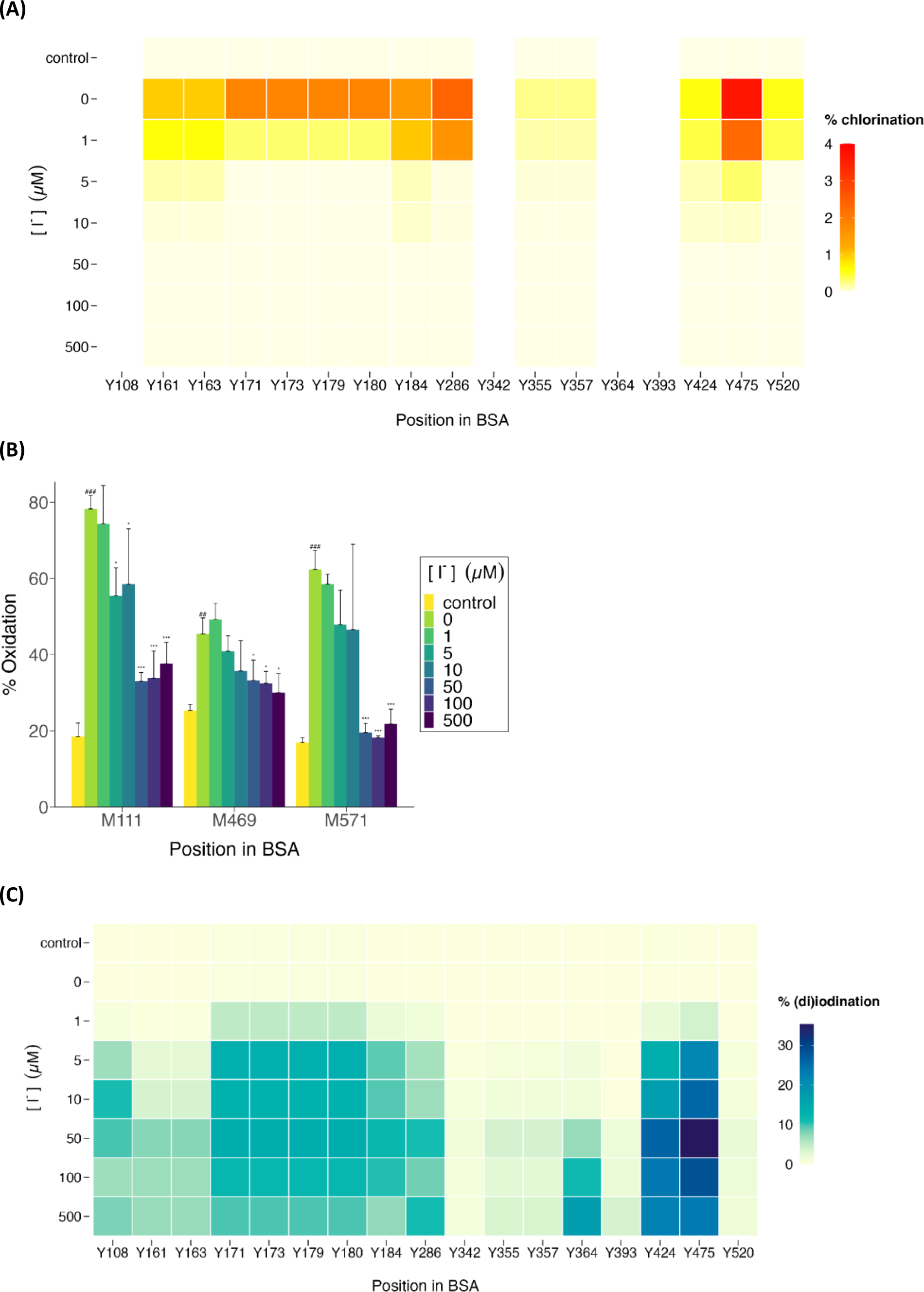
Mapping of protein modifications on bovine serum albumin induced by MPO/H_2_O_2_/Cl⁻ systems in the absence or presence of increasing concentrations of I⁻. Bovine serum albumin (2 µM) was treated with MPO (0.1 µM), H_2_O_2_ (20 µM, 10-fold molar excess over the BSA concentration), and Cl⁻ (100 mM) in the absence or presence of increasing I⁻ concentrations (1 - 500 µM) for 2 h at 37 °C before reduction/alkylation, trypsin digestion and LC-MS/MS peptide mapping. (A) Heat map of the extent of chlorination at all Tyr residues (top row; except Y54, Y376 which were detected, but not identified as being modified; and Y434 which was not covered in the peptides detected), and the effect of increasing I⁻ (subsequent rows, with increasing concentrations down the map). (B) Extent of oxidation at each Met residue (with the exception M208 which was detected in a peptide with a weak signal, but not identified as being modified) on BSA by the above oxidation systems in the absence of I⁻, MPO and H_2_O_2_ (bars labelled ‘control’) and in the presence of MPO, H_2_O_2_ and increasing I⁻ as indicated. (C) Heat map of percentage iodination/di-iodination at all Tyr residues (except Y54, Y376 which were not identified as being modified; and Y434 which was not covered in the peptides detected) in the absence of any I⁻ (top row, labelled ‘control’) and in the presence of increasing I⁻ (subsequent rows). Data are presented as mean + SD from 3 independent experiments. # Indicates a statistical difference between the control against the oxidized sample in the absence of I⁻ (0 µM I⁻) as determined by two-sample t-test. * Indicates statistical significance against the oxidized sample in the absence of I^-^ (0 µM I^-^) as determined by one-way ANOVA with post hoc Dunnett’s test. * *p* < 0.05, ** *p* < 0.01, *** *p* < 0.001.

Met oxidation was detected on 6 out of the 7 residues in MSA, with 5 of these validated and quantified based on peak shape, retention time, mass error, isotopic distribution, and MSMS library match. Control samples of MSA contained significant amounts of oxidized Met (Supplementary Figure 4, 5 - 25% occupancy); this is likely to arise, at least in part from the biological origin of this material and oxidation during purification and processing. However, the extent of oxidation at 3 of these residues (M159, M322, M572) increased significantly on treatment with LPO/H₂O₂/I⁻ systems containing increasing levels of I⁻, though to variable extents (Supplementary Figure 4), consistent with a degree of selectivity in oxidation of this residue. Whilst the extent of oxidation at M322 and M572 increased with I⁻ concentrations over the range 1 - 10 µM, this was reversed at high I⁻, possibly due to the formation of alternative products (e.g. the di-oxygenated sulfone [36], which was not quantified) that distort the % occupancy data. These two residues were the most sensitive to modification reaching a maximum of 62% and 26% oxidation.

Modification (mono- and di-oxidation) was also detected at W238, with this showing a similar pattern of behaviour to the Met residues. Limited Trp oxidation was detected in the control samples, probably for the same reasons as outlined above; but the extent of modification was increased on treatment with LPO/H₂O₂/I⁻ systems containing 5 - 10 µM I⁻ (Supplementary Table 1).

### 3.3. Increasing concentrations of iodide modulate the modifications induced on anastellin and serum albumins by myeloperoxidase systems

In the light of the above data indicating facile iodination of Tyr residues on both anastellin and MSA induced by enzymatically generated HOI, experiments were performed to examine whether iodination also occurred when I⁻ and Cl^-^ were present as competing substrates using a MPO system. Thus the proteins were treated with the MPO/H₂O₂/Cl⁻ ± I⁻ system, with variable amounts of I⁻ and constant concentrations of MPO, H₂O₂ and Cl⁻, reflecting pathophysiological conditions where the concentration of Cl⁻ is likely to be high (100 – 150 mM) and essentially constant.

For anastellin, the protein was exposed to MPO (100 nM) / H₂O₂ (50 μM) / Cl⁻ (100 mM) in both the absence and presence of I⁻ (at 10, 100 or 1000 μM) for 2 h at 37°C. The protein samples were then digested with trypsin and Glu-C and subjected to peptide mass mapping. Peptides corresponding to >95 % of the protein sequence were detected (see also [24]), and all 4 Tyr residues present in anastellin were detected. In the presence of the complete MPO system, but with no added I⁻, loss of the parent Tyr residues was detected with the extent of loss being dependent on the Tyr residue location, with this ranging from ∼6% to 45% (Supplementary Table 2). This loss was accompanied by the detection of ions assigned to 3-chlorotyrosine (3-ClTyr; detected as *m/z* +34 Da mass shifts) and also limited amounts of the dichlorinated species 3,5-di-ClTyr (detected as *m/z* +67.9 Da mass shifts). In the presence of I⁻ as well as Cl⁻, significant decreases in the yield of the chlorinated species were observed (Supplementary Table 2), with concomitant detection of ions assigned to both mono- and di-iodinated Tyr residues (3-iodoTyr and 3,5-di-iodoTyr, mass shifts of +125.9 and +251.8 respectively; Supplementary Table 2). The decrease in chlorination and increase in iodination occurred in a dose-dependent manner with increasing I⁻ concentrations in the reaction mixture. Furthermore, in the presence of I⁻ (at 10, 100 or 1000 μM) evidence for the formation of both mono- and di-iodinated His products were detected for the His residue present in the peptide GNAPQPSHISK (Supplementary Table 2). The yield of iodinated His containing peptides increased with the I⁻ concentration, and accounted for ∼25% of the parent peptide intensity at the highest I⁻ concentration examined. No evidence was obtained for the formation of significant levels of chlorination at the same site (Supplementary Table 2). His residues present on the His-tag are also likely targets of modification, but further analysis was precluded due to the presence of complex, overlapping isomeric species with poor fragmentation spectra. Nevertheless, these data are consistent with I⁻ acting as a competitive substrate for the MPO system, and the formation of both iodinated Tyr and His products helps account for the high levels of iodine incorporation detected in the intact protein experiments (see above).

Analogous experiments were carried out for the serum albumins using BSA, as previous data on the effects of enzymatic MPO systems and reagent HOCl, on serum albumins [37, 38], has been carried out with this protein rather than MSA. BSA and MSA have a very high extent of sequence identity and homology (69.7% sequence similarity, 19 homologous Tyr, 3 non-homologous Tyr in MSA, 2 non-homologous Tyr in BSA; Supplementary Fig. 6). Initial studies examined the basal condition of modification of BSA by MPO/H₂O₂/Cl⁻, in the absence of added I⁻, using the same peptide mass mapping approach described above. Under the reaction conditions employed (100 nM MPO, 3 μM BSA, 30 μM H_2_O_2_ added as a single bolus, 100 mM Cl⁻; incubation at 37 °C for 2 h), both oxidation (at Met) and chlorination (at Tyr) were detected (Fig. 3A,B). Modification at other residues were either not quantified (e.g. at Cys34, though this is a known target for HOCl [38]) or not detected (modifications at Trp, His, Lys, cystines) due to computational limitations arising from searching for large numbers of modifications in open searches. Chlorination (to give 3-ClTyr) was detected to significant levels at multiple Tyr residues, including Y161, Y163, Y171, Y173, Y179, Y180, Y184, Y286 and Y475. Chlorination at Y475 was particularly marked with this occurring to a maximum extent of ∼4 %. These chlorinated species were not detected when any component of the complete reaction system was absent. In contrast, the complete reaction system induced significant levels of oxidation at 3 out of 4 Met residues present in BSA, though to varying extents (the fourth Met residue, M208, was detected in a peptide with a weak signal and the corresponding oxidised peptide was not detected). As with the MSA experiments described above, some oxidation was detected in the parent (untreated) BSA samples, though the extent of modification was enhanced by treatment with the enzyme system. The extent of modification at some sites was very high, with, for example, the % modification at M111 reaching 78.3 %, and high values also detected for M469 (49.2 %) and M571 (62.4 %) (Fig. 3B).

The addition of low concentrations of I⁻ to the above MPO reaction systems caused dramatic changes in both the pattern and extent of protein modification, with ions assigned to iodinated Tyr residues detected (Fig. 3C). The detection of these species was accompanied by a loss of 3-ClTyr (Fig. 3A), consistent with a switch from chlorination to iodination, with this occurring in a dose-dependent manner with increasing amounts of I⁻ in the reaction mixture. This changeover from chlorination to iodination occurred even with 1 μM I⁻ (Fig. 3A,B). For the most heavily modified Tyr residues (Y424, Y475) ions were also detected consistent with the presence of 3,5-di-iodoTyr (*m/z* +252 Da mass shifts). Detection of these di-iodo species occurred with modest levels of I⁻ (≥ 10 μM), with the % of di-iodination (occupancy) being ∼30-35% at I⁻ concentrations above 10 µM. This is in marked contrast to the extent of 3-ClTyr formation obtained with MPO-derived HOCl, with the extent of modification being ∼10-fold higher for iodination than chlorination (cf. color coded extents of modification in Fig. 3A,C). With I⁻ concentrations in the 1 - 5 µM range approximately equal yields of 3-ClTyr and 3-iodoTyr were detected, despite the ∼10,000-fold difference in the concentrations of I⁻ versus Cl⁻. In a similar manner to the reduction in the yield of 3-ClTyr detected with increasing I⁻ concentrations, a significant decrease in Met oxidation was also detected at high I⁻, with the % modification values for Met decreasing to near control levels with ≥ 10 µM I⁻ (Fig. 3B).

These data indicate that there are major differences in the selectivity of HOCl versus HOI, with the preference for modification at the thioether (R-S-R) groups of Met being much lower for HOI than HOCl (see also Discussion).

### 3.4. Structural analysis by SDS-PAGE indicates modest effects of HOI, but more major effects of HOCl with this modulated by increasing concentrations of iodide

SDS-PAGE analysis of both mouse and human serum albumins treated with LPO/H₂O₂/I⁻ resulted in the detection of minimal gross structural changes (data for MSA presented in Supplementary Fig. 5). Even very high concentration of both H₂O₂ and I⁻ (> 300 µM and 10 mM, respectively) yielded only a modest broadening of the parent albumin band (Supplementary Fig. 5). No evidence of additional protein dimers/cross-links was obtained above that already present in the commercial protein samples, and only low levels of protein cleavage (bands of lower molecular mass than the parent protein) were detected.

In contrast to the above, the MPO/H₂O₂/Cl⁻ system (i.e. formation of HOCl) gave rise to significant structural damage to serum albumins when high concentrations of H₂O₂ were employed, with both protein aggregates and cleavage detected by SDS-PAGE (Fig. 4). These observations are consistent with previous studies [37, 38]. With moderate excesses of H₂O₂ (25-fold relative to protein), the parent band at ∼60 kDa remained mostly intact, with modest concentrations of species with lower molecular mass detected after HOCl treatment (Fig. 4). The presence of increasing concentrations of I⁻ in the MPO/H₂O₂/Cl⁻ system resulted in less intense bands from the cleavage products (Fig. 4).

**Figure 4.**
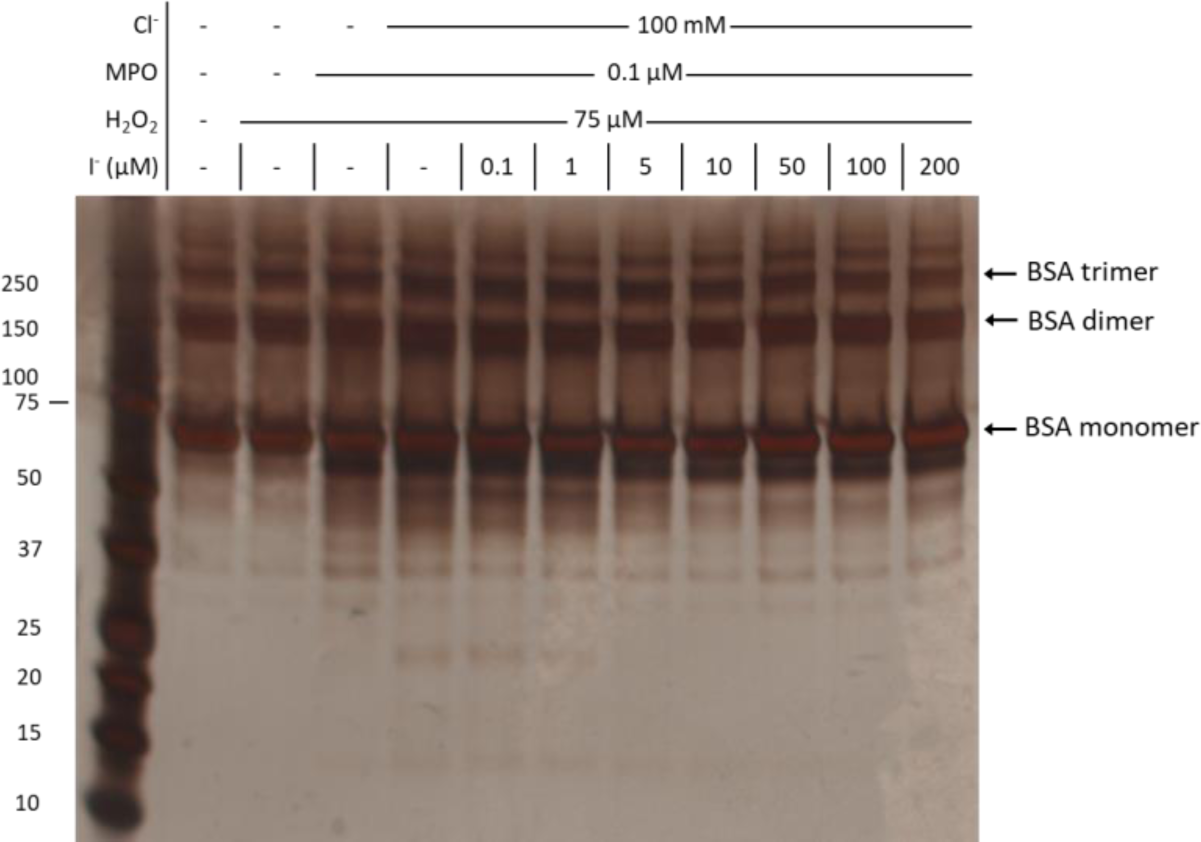
I⁻ reduces structural changes induced by MPO/H_2_O_2_/Cl⁻ system to bovine serum albumin. Bovine serum albumin (3 µM) was treated with MPO (0.1 µM), H_2_O_2_ (25-fold molar excess, 75 µM), and Cl⁻ (100 mM) in the absence or presence of increasing I⁻ (0.1 - 200 µM) for 2 h at 37 °C in chelexed phosphate buffer, pH 7.4. Protein was detected by silver staining following SDS-PAGE separation under reducing conditions. The molecular masses of the markers present in the first lane are indicated at the left-hand side of the figure. The position of the bands assigned to BSA monomer, dimer and trimer are indicated on the right-hand side of the panel.

In a similar manner to the serum albumin systems described above, treatment of anastellin (which runs as a band at ∼12 kDa, which is slightly higher than the molecular mass for this recombinant protein of 9.4 kDa, but consistent with previous reports [24]) with LPO/H₂O₂/I⁻ resulted in minimal gross structural changes to the parent protein as detected by SDS-PAGE. No evidence was detected for dimers or higher oligomers, or low molecular mass fragments. In contrast, treatment with MPO/H₂O₂/Cl⁻, in the absence of I⁻, resulted in the detection of dimeric species (band at ∼20 kDa in Fig. 5; the higher mass band on these gels is from the heavy chain of MPO). These dimers were not detected with untreated anastellin, or anastellin treated with H₂O₂ in the absence of MPO. Inclusion of I⁻ at increasing concentrations, decreased the dimer yield in a dose-dependent manner (Fig. 5), such that with ≥ 50 µM I⁻ very little dimer was detected.

**Figure 5.**
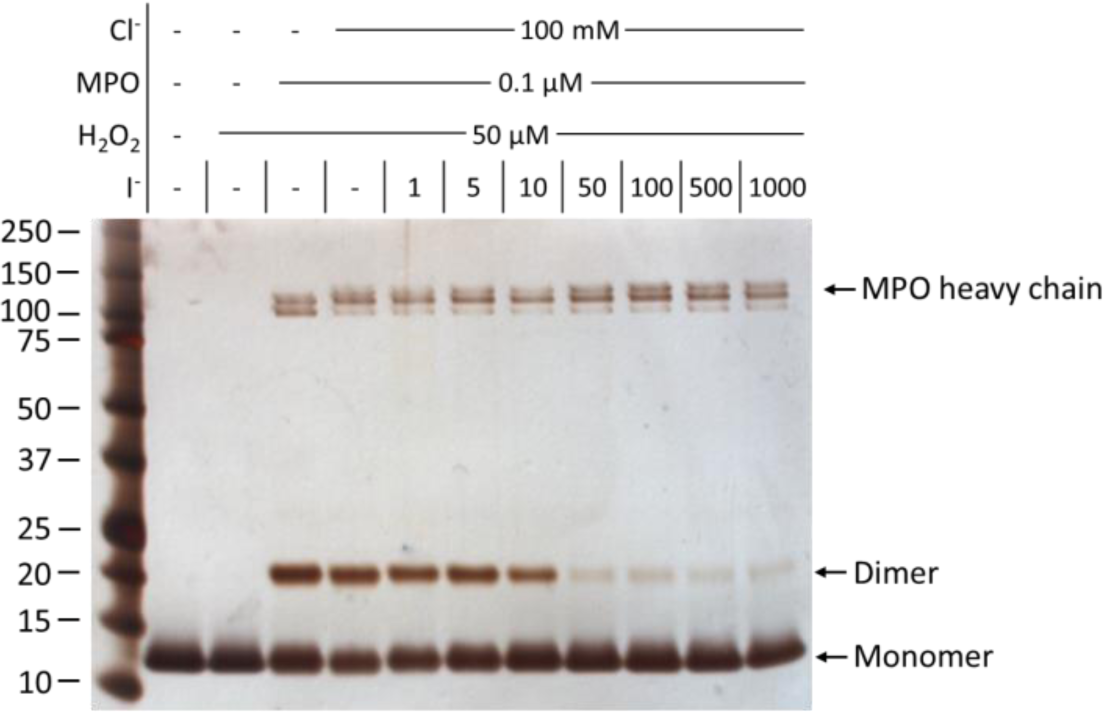
The formation of anastellin dimers induced by MPO/H_2_O_2_/Cl⁻ is inhibited by increasing concentrations of I⁻. Anastellin (5 µM) was treated with MPO (0.1 µM), H_2_O_2_ (50 µM, 10-fold molar excess over anastellin), and Cl⁻ (100 mM) in the absence or presence of increasing I⁻ (1 - 1000 µM) for 2 h at 37 °C in chelexed phosphate buffer, pH 7.4. Protein was detected by silver staining following SDS-PAGE separation under non-reducing conditions. The molecular masses of the markers present in the first lane are indicated at the left-hand side of the figure. The position of the bands assigned to anastellin monomer and dimer are indicated on the right-hand side of the gel image.

These data are consistent with HOI inducing limited or no inter-chain cross-linking, or fragmentation of the peptide backbone. In contrast, HOCl induces both types of structural modifications, and particularly cross-linking with anastellin. However, when I⁻ is present in the MPO systems, competition occurs between I⁻ and Cl⁻ as substrates for MPO, with the products from oxidation of I⁻ (i.e. HOI) inducing lower levels of gross structural alterations than HOCl.

## 4. Discussion

With the exception of the reactions of thyroid peroxidase, the role of peroxidase-mediated iodination of proteins and the effects of HOI on mammalian systems have been poorly explored. This may be due to, at least in part, the low concentrations of I⁻ present in plasma [34, 35], and therefore the assumption that this anion is of limited relevance to the total oxidant yield and protein modification. However there is now growing interest in the role of I⁻ as a peroxidase-substrate, and HOI as a biologically-relevant oxidant, with the lack of information on the role of these species being a key knowledge gap.

The increasing interest is this area, has arisen from reports that I⁻ can reduce HOCl-induced oxidative damage derived from MPO in the presence of H₂O₂ [21], and the detection of iodinated proteins from systems containing activated human neutrophils, in sputum from people with cystic fibrosis and in skin abscesses from people infected with *S. aureus* [23]. In addition, I⁻ supplementation has been reported to afford protection against ischemia-reperfusion injury in a number of animal models [12–14], though the origin of this effect has been ascribed to metabolic changes, rather than effects on peroxidase-mediated reactions. These may however be interlinked as thyroid hormone concentrations are dependent on I⁻ concentrations and the activity of thyroid peroxidase. Furthermore, in the case of the neutrophil-mediated modifications, the resulting products appear to have distinct stimulatory effects on chemotaxis and cytokine release [23]. These data indicate that protein iodination may be more common than previously appreciated (possibly as a result of elevated concentrations of I⁻ within neutrophil phagolysosomes [23]) or other factors (see below), and be of biological importance.

In the current study, treatment of multiple model proteins with LPO/H₂O₂/I⁻ (i.e. in the absence of other halide/pseudohalide substrates) has been shown to give rise to efficient incorporation of iodine into the proteins. This appears to be very facile, with high quantitative conversion of the initial iodinating species (presumed to be HOI or I^+^) into iodinated amino acids on the target protein. This is consistent with previous data, where this enzyme system has been used to carry out radio-labelling of proteins [39]. The data obtained for anastellin by direct intact-protein LC-MS, has provided evidence for the incorporation of large numbers of iodine atoms (up to 20). This number far exceeds the number of Tyr residues (4) present in the protein. Thus even if di-iodination occurs at each of these residues (to give initially 3-iodoTyr and subsequently 3,5-di-iodoTyr), this maximally accounts for 8 iodine atoms. These data are therefore consistent with iodination at other sites, which are presumed to be on the two Trp, and nine His residues (3 in the parent protein sequence and 6 in the His tag). The peptide mass mapping studies carried out on this protein have provided for both mono- and di-iodination at one of the His resides, consistent with this hypothesis. Interestingly, no significant chlorination was detected at this same residue when treated with the MPO/H_2_O_2_/Cl⁻ system (Supplementary Table 2) possibly for kinetic reasons (i.e. much more rapid iodination than chlorination). The generation of iodinated Trp residues is consistent with the previous detection of corresponding mono- and di-chlorinated species from this amino acid side chain [40], though the iodinated species appear to be generated in higher yield. In molar terms the LPO/H₂O₂/I⁻ systems appears to be extremely efficient, with 50 µM of H_2_O_2_ yielding ∼30 µM of iodinated products on MSA (i.e. ∼60% yield), whereas the MPO/H₂O₂/Cl⁻ ± I⁻ system is less efficient, giving ∼5 µM of products from 20 µM of oxidant (∼25% yield) (Fig. 7).

In contrast to the detection of multiple iodinated residues with anastellin, with MSA and BSA most of the iodination appears to occur at Tyr with this giving both 3-iodoTyr and 3,5-di-iodoTyr. There are multiple potential reasons for this difference. One is that some of the iodinated species decompose during processing and are therefore not detected. This could not be easily tested as the spectra obtained from the intact MS experiments with serum albumins could not be mass-resolved (Supplementary Figure 1).

However, it is also possible that this difference arises from the lower relative abundance of Trp and His in albumins (2 Trp, 17 His and 20 Tyr residues in BSA; 1 Trp, 17 His and 22 Tyr in MSA) when compared to anastellin (2 Trp, 9 His and 4 Tyr), and a greater surface exposure of the Trp residues in anastellin, compared to the (mostly) buried Trp residues in MSA and BSA (W238 in MSA; W158 and W237 in BSA; Fig. 6). Oxidation was also detected at the Met residues present in these serum albumins (7 in MSA and 4 in BSA). This was not observed with anastellin as it lacks any Met.

**Figure 6.**
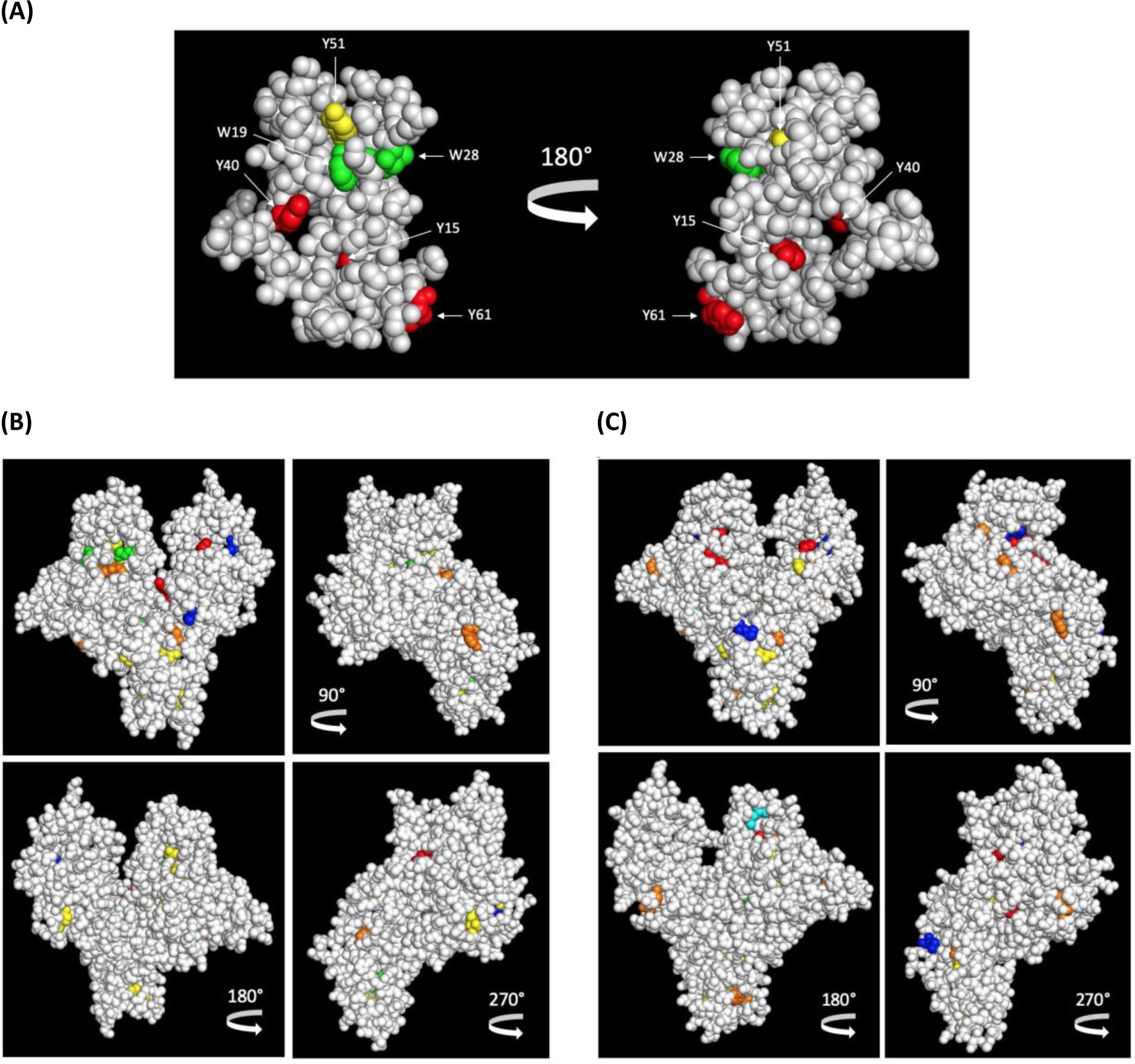
Structure of anastellin (A) and serum albumins (B, C) showing position of all Tyr, Trp and (for serum albumins) Met residues (van de Waals radius). Residues are color coded to indicate the extent of modification by the oxidant systems. Tyr residues are colored based on their % (di)iodination (yellow < orange < red) while Met and Trp residues are colored according to % oxidation (cyan < blue). Green = no modification. (A) Anastellin (PDB entry: 1Q38) modified by MPO/H_2_O_2_/Cl^-^/I⁻ system. Tyr; red: ∼80% iodination, yellow: < 20% iodination. Extent of modification is based on Supplementary Table 2. (B) BSA (PDB entry: 4F5S) modified by MPO/H_2_O_2_/Cl^-^/I⁻ system. Extent of modification is based on Fig. 3B and 3C. Tyr; red: > 20% iodination, orange: > 10% iodination, yellow: < 10% iodination. Dark grey = not covered.

**Figure 7.**
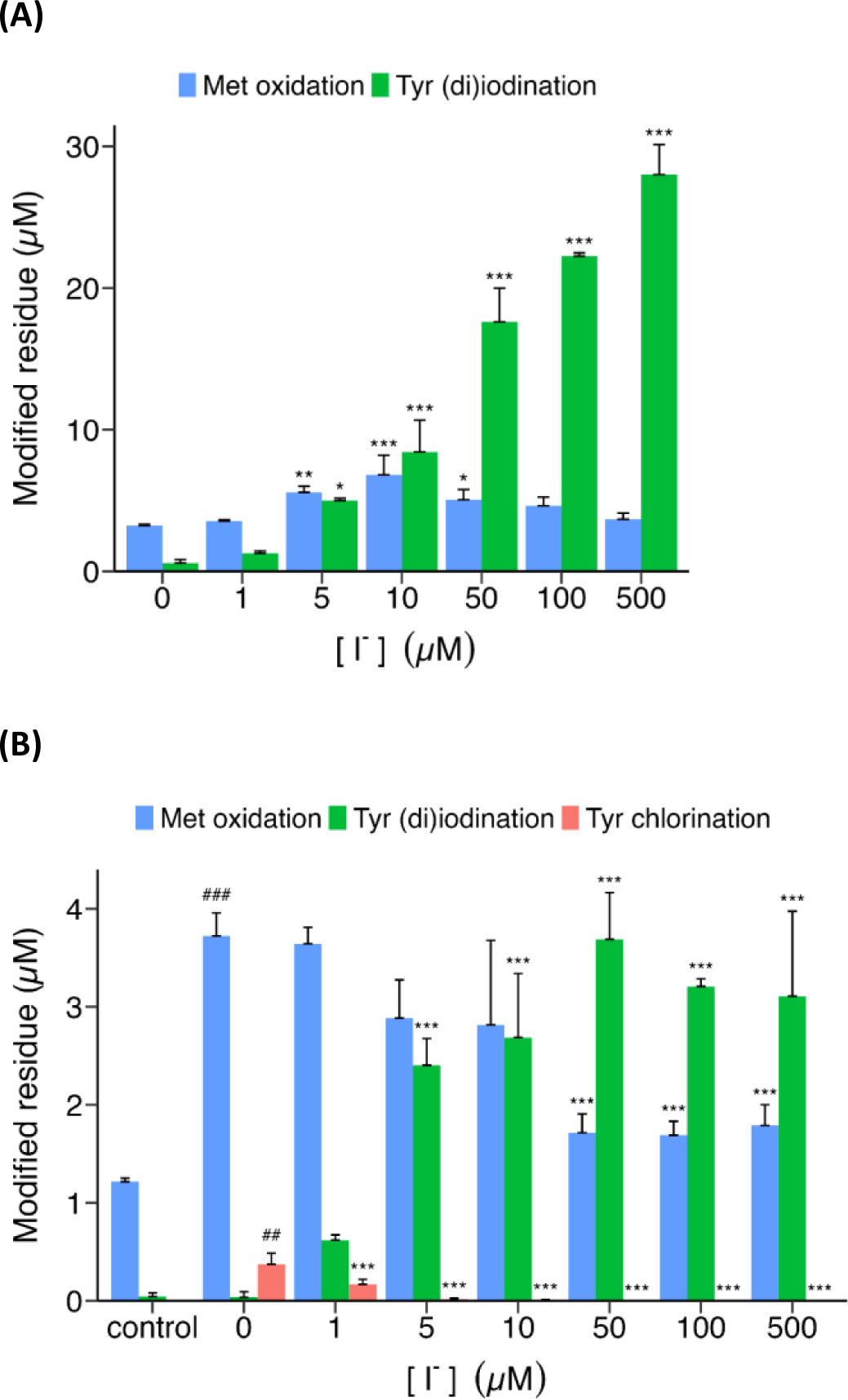
Comparison of residue consumption by HOCl versus HOI and extent of formation of products at Met versus Tyr residues. (A) Sum of the extents of formation of products at Met and Tyr residues on MSA (5 µM) treated with LPO (1.5 µM) and H_2_O_2_ (50 µM, 10-fold molar excess over MSA) in the absence or presence of increasing I⁻ concentrations (1 - 500 µM) for 2 h at 37 °C before reduction/alkylation, digestion and LC-MS/MS peptide mapping. The resulting extents of modification at each residue type (Met – oxidation; Tyr – iodination) were then summed. Data replotted from Fig. 2C and Supplementary Fig. 4. (B) Sum of the extents of formation of products at Met and Tyr residues on BSA (2 µM) treated with MPO (0.1 µM), H_2_O_2_ (20 µM, 10-fold molar excess over BSA), and Cl⁻ (100 mM) in the absence or presence of increasing I⁻ (1 - 500 µM) for 2 h at 37 °C before reduction/alkylation, trypsin digestion and LC-MS/MS peptide mapping. The resulting extents of modification at each residue type (Met – oxidation; Tyr – chlorination or iodination) were then summed. Data replotted from Fig. 3. Data are presented as mean + SD from 3 independent experiments. * Indicates statistical significance against the oxidized sample in the absence of I^-^ (0 µM I^-^) as determined by one-way ANOVA with post hoc Dunnett’s test. # Indicates statistical significance of the control against the oxidized sample in the absence of I^-^ (0 µM I^-^) as determined by two-sample t-test. * *p* < 0.05, ** *p* < 0.01, *** *p* < 0.001.

Met; blue = oxidation. (C) HSA (PDB entry: 1AO6) modified by LPO/H_2_O_2_/I⁻ system. The crystal structure of MSA is not determined so the structure of HSA was used in place. The extent of modification is based on Fig. 2B, 2C and Supplementary Fig. 4. Tyr; red: > 20% iodination + significant loss of parent ion, orange: > 20% iodination, yellow: < 20% iodination. Met/Trp; blue = significant oxidation, cyan = non-significant oxidation. PyMOL (version 2.5.2) was used to make the figure.

Whilst these data indicate that HOI (or I^+^) is a highly efficient agent with regards to iodination of Tyr and His, the use of LPO/H₂O₂/I⁻ does not allow analysis of the relative contributions of iodination versus chlorination. This was therefore examined with a MPO/H₂O₂/Cl⁻ system in the presence of I⁻ over a wide range of I⁻ concentrations spanning from normal physiological levels through to supraphysiological (see below). The experiments with both anastellin and serum albumins, show that MPO/H₂O₂/Cl⁻/I⁻ systems are also effective iodinating systems when the concentration of I⁻ is > 1 μM. In the *absence* of I⁻, chlorination at specific Tyr, and Met oxidation was detected, consistent with previous data (e.g. [9]). However iodination at a large number of the total complement of Tyr residues was detected with low μM concentrations of I⁻. Both mono- and di-iodination was detected at multiple residues, with a concomitant loss of the chlorinated Tyr and His species, and also a decrease in the extent of Met oxidation. The latter observation suggests that Tyr and His iodination is competitive with Met oxidation even allowing for the lower number of Met residues compared to Tyr in these proteins. This is the very different to the situation with HOCl, where the rate constant for Met oxidation is ∼10^5^-fold greater than for Tyr chlorination [41, 42], and indicates that the rate constants for iodination of Tyr and His by HOI are likely to be in the same range as for Met oxidation by HOCl [41, 42].

Mapping of the sites of modification induced by the MPO/H₂O₂/Cl⁻ ± I⁻ systems to the 3-dimensional structure of HSA (Fig. 6) shows that the modified Tyr residues are widely spread over the structure, though these are confined to one face of the protein. These are color-coded in Fig. 6 to the extent of modification detected. No structures of MSA are available, though sequence alignment of murine, bovine and human serum albumin reveals a high amino acid similarity (Supplementary Fig. 6 [43]) and the sequence differences between these homologs do not appear to change the protein tertiary structure. This wide distribution is consistent with the generation of a diffusible oxidant, such as HOI, rather than oxidation via an enzyme-bound intermediate. Whether the distribution is influenced by possible binding of LPO or MPO to the serum albumins is unclear, though evidence has been reported for the binding of MPO to several proteins (reviewed [36]), including albumins [44], though the binding sites on albumins are unknown. Such binding has been reported, in some cases, to enhance enzymatic activity [45].

Unfortunately, the reaction rate constants for HOI towards specific amino acids are unknown. This work suggests that HOI is dramatically more reactive towards Tyr than HOCl, and that the reaction of HOI with Met is correspondingly less competitive. This is exemplified by the data in Fig. 7, which indicates that the extent of Met loss, and Met sulfoxide formation, are low and remain relatively constant for the LPO/H₂O₂/I⁻ system (with low extents of Met sulfoxide detected in the control protein samples) with increasing I⁻ concentrations, whereas the yield of Tyr iodination products increases dramatically (Fig. 7A). In contrast for the MPO/H₂O₂/Cl⁻/I⁻ systems, the yield of Met sulfoxide and also chlorinated Tyr residues increases significantly when the protein is exposed to the intact MPO/H₂O₂/Cl⁻ system (with no I⁻) when compared to the control samples (compare first two sets of bars in Fig. 7B). However with increasing I⁻ present in the MPO system, and consequent competition between the two anions, the yields of both Met sulfoxide and Tyr chlorination decrease significantly and the yield of iodinated Tyr (and His for anastellin) increases dramatically (Fig. 7B). Tyr chlorination is eliminated in this system by addition of 1 – 10 µM I⁻ (Fig. 3A and Supplementary Table 2). HOBr is less reactive towards Met than HOCl [41, 46], and it is likely that HOI will follow this trend due to the electronegativity of the iodine atom (I > Br > Cl). Thus both HOBr and HOI react ∼3 orders of magnitude faster towards phenols (to give the corresponding halogenated species) than HOCl [47–50].

The switch from chlorination to iodination occurs at a much lower concentration of added I⁻ than would be predicted from the respective rate constants for reaction of the two halide ions with Compound I of MPO, with these values differing by a factor of 288 (see above). Thus, with the 100 mM Cl⁻ employed in these experiments it would be expected that ∼ 0.35 mM I⁻ would be required to give approximately equal concentrations of HOCl and HOI, if competitive reaction with Compound I was the only process involved. It is known, however, that HOCl can also rapidly oxidise I⁻ to HOI in a direct molecular reaction, with this occurring with a rate constant, *k*, of ∼1.4 x 10^8^ M^-1^ s^-1^ [51]. It is therefore likely that some of the HOCl formed by MPO is converted by this reaction to HOI, thereby diminishing chlorination and enhancing iodination [52]. Whilst this process may be of importance *in vitro*, it may be less significance *in vivo* (or other complex systems), due to the abundance of other targets for HOCl [42].

Previous studies have variously postulated the involvement of HOI, I^+^ and I^•^ in the generation of iodinated Tyr residues, and photochemical generation of I^•^ has been shown to give rise to protein iodination [53]. However it is likely that HOI or I^+^ are the species involved in the reactions studied here, rather than I^•^, as the redox potential needed for oxidation of I⁻ to I^•^ is greater than that for the Compound I to Compound II, or Compound II to ferric state, reactions of the MPO peroxidatic cycle, making radical formation unfavourable.

Although both iodination systems appear to give significant levels of iodinated Tyr, HOI does not appear to result in marked gross structural changes on the targeted proteins, at least as detected by SDS-PAGE. Whilst no significant changes were detected for the serum albumins with the LPO/H₂O₂/I⁻ system, the MPO/H₂O₂/Cl⁻ system induced significant anastellin dimerization, and modest aggregation and fragmentation with the serum albumins. These effects of HOCl are consistent with previous data [37].

While the precise mechanism of HOCl-derived anastellin cross-linking remain elusive (as this protein does not contain Cys or Met), it may involve Tyr-Tyr, Tyr-Trp, or Trp-Trp dimers generated by TyrO^•^ or TrpN^•^ radicals [24, 54]. Addition of I⁻ into the MPO/H_2_O_2_/Cl⁻ system inhibited the dimerization of anastellin (Fig. 5). This is consistent with previous data with fibronectin where a protective effect of I⁻ against dimerization was observed [11]. Furthermore, the absence of dimer formation induced by LPO/H₂O₂/I⁻ (and inhibition of MPO-mediated oligomerization) is consistent with an absence of I^•^, as many cross-linking processes involve radical-radical reactions [55].

Although the SDS-PAGE experiments indicated an absence of gross structural changes generated by HOI, the intact LC-MS data obtained with anastellin indicates that iodination of this protein increases the protein hydrophobicity. This was manifested as an increased retention of the modified protein on the LC column (Fig. 1B). This effect is consistent with calculated LogP values (partition coefficient of a solute between octanol and water, at near infinite dilution) of −0.56 vs. −1.50 for Tyr vs 3-iodoTyr, respectively (determined using the ChemAxon program ‘Chemicalize’, https://chemicalize.com/). These data are consistent with that reported for chlorinated- and iodinated α-defensin proteins [23]. The altered surface properties, and greater hydrophobicity induced by Tyr iodination may give rise to altered interactions with partner proteins including receptors, and therefore explain (at least in part) the altered biological properties of the iodinated (and to a lesser extent chlorinated) proteins and peptides (cf. data in [23]).

These results suggest that protein iodination (external to the thyroid) may be particularly relevant in situations of high peroxidase expression and / or high I⁻ concentrations. MPO expression is known to vary between individuals due to genetic factors and inflammatory status, with this enhancing the risk of disease [56–60]. Elevated levels of other peroxidases, including LPO and eosinophil peroxidase have been reported in various diseases (e.g. [61, 62]). Whilst the levels of I⁻ in plasma or serum are typically 0.1 – 1 µM [34, 35, 63], these values vary widely, with diet being a major factor [64]. Values in other biological fluids can be markedly higher due to selective concentration or transport. Concentrations in human breast milk are ∼2 µM [65], where the LPO/H_2_O_2_/I⁻ system has antimicrobial properties, with this being linked to HOI formation [66]. Elevated I⁻ levels are also present in saliva, both in healthy subjects (mean values ∼ 1 µM [65]), after subject exposure to iodinated contrast agents used in coronary angiography (mean values 16 vs. 332 µM, with the highest values being ∼1500 µM [65]), and as a result of treatment with the antiarrhythmic drug amiodarone (a 3,5-di-iodotyrosine derivative; saliva values up to ∼100 µM [67]). Serum I⁻ levels are also enhanced by iodinated contrast agent exposure (mean concentrations ∼ 1.5 µM, peak values ∼ 7 µM [65, 68]). Airway mucosa and secretions also have significantly elevated I⁻ levels compared to serum levels [69].

The concentration of I⁻ within cells, apart from those in the thyroid, is less well researched, but elevation, relative to plasma / serum values, may occur via the activity of ion transporters or channels, including the Na^+^/I⁻ symporter (NIS; expressed in multiple cells [70]), the cystic fibrosis transmembrane conductance regulator (CFTR), anotamin-1 (ANO1, also known as TMEM16A) or the anion exchanger pendrin (SLC26A4) [65]. The action of these transporter/channels (but not NIS, as neutrophils do not express this protein) may underlie the detection of iodinated proteins within neutrophil phagolysosomes [23].

Together these data suggest that peroxidase-mediated generation of HOI, and formation of iodinated species, may be relevant in individuals with acute or chronic inflammation, in those with high dietary iodine/iodide/iodinated compound consumption, or following exposure to treatments that elevate I⁻ levels. Data indicate that deliberate elevation of I⁻ is protective in humans in cases of radiation exposure (where oral I⁻ tablets are employed to prevent radioactive I⁻ accumulation), to prevent ischemic-reperfusion injury [12–14], as a treatment for COVID and other viral diseases (povidone-iodine [71] or I⁻ salts [72]) and sepsis [73].

## Acknowledgements

The authors are grateful for financial support from the Novo Nordisk Foundation (NNF20SA0064214 to M.J.D.), the Lundbeck Foundation (R322-2019-2337 to L.F.G.) and the Carlsberg Foundation for an infrastructure grant (CF19-0451 to P.M.H.).

## Abbreviations

ACN: acetonitrile
3-ClTyr: 3-chlorotyrosine
3,5-di-ITyr: 3,5-di-iodotyrosine
ESI: electrospray ionization
FA: formic acid
HOCl: the physiological mixture of hypochlorous acid and its conjugate anion ^−^OCl
HOI: the physiological mixture of hypoiodous acid and its conjugate anion ^−^OI
HPLC: high-performance liquid chromatography
3-ITyr: 3-iodotyrosine
LC-MS: liquid chromatography-mass spectrometry
LPO: lactoperoxidase
MPO: myeloperoxidase
MS: mass spectrometry
PTM: post-translational modification
TFA: trifluoroacetic acid
UPLC: ultra high-performance liquid chromatography.

