## Supplemental Figures for "Elevated levels of iodide promote peroxidase-mediated protein iodination and inhibit protein chlorination"

**SUPPLEMENTARY DATA**

(A)
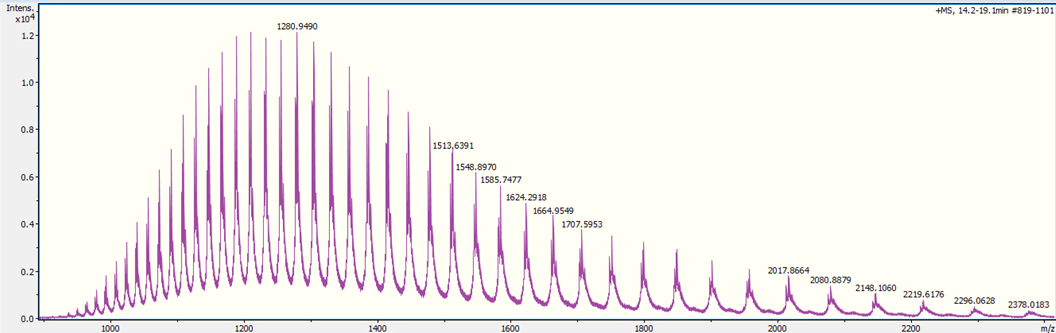

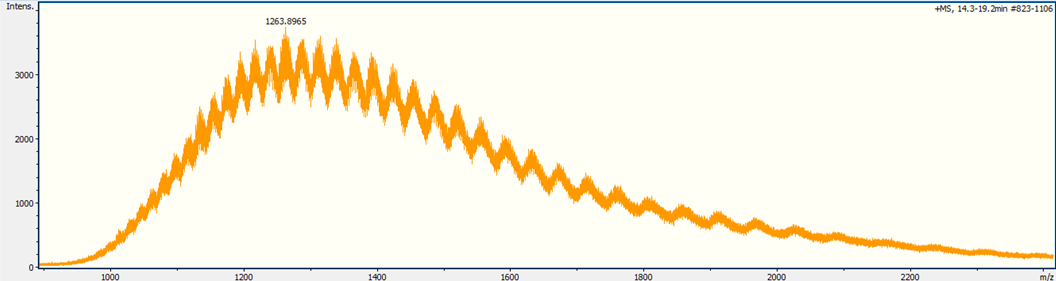
**(B)**

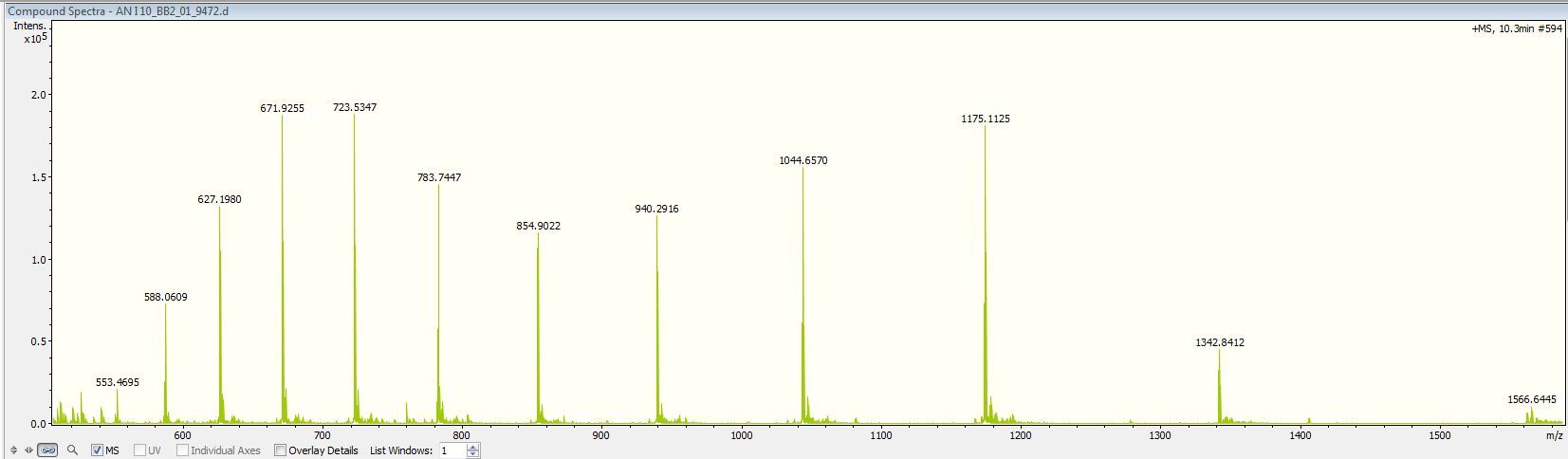
**(C)**

**Supplementary Figure 1.** LC-MS intact protein analysis of human serum albumin (HSA, 3 µM) treated with LPO (1.5 µM), H_2_O_2_ (10-fold molar excess, 30 µM), and I⁻ (1000 µM) for 2 h at 37 °C in chelexed phosphate buffer, pH 5.8. Mass spectrum of: (A) untreated and (B) oxidant-treated HSA; (C) untreated anastellin. The different species for oxidant-treated albumin could not be resolved and quantified due to the large number of charge states, while for anastellin the charge states were extremely well resolved.

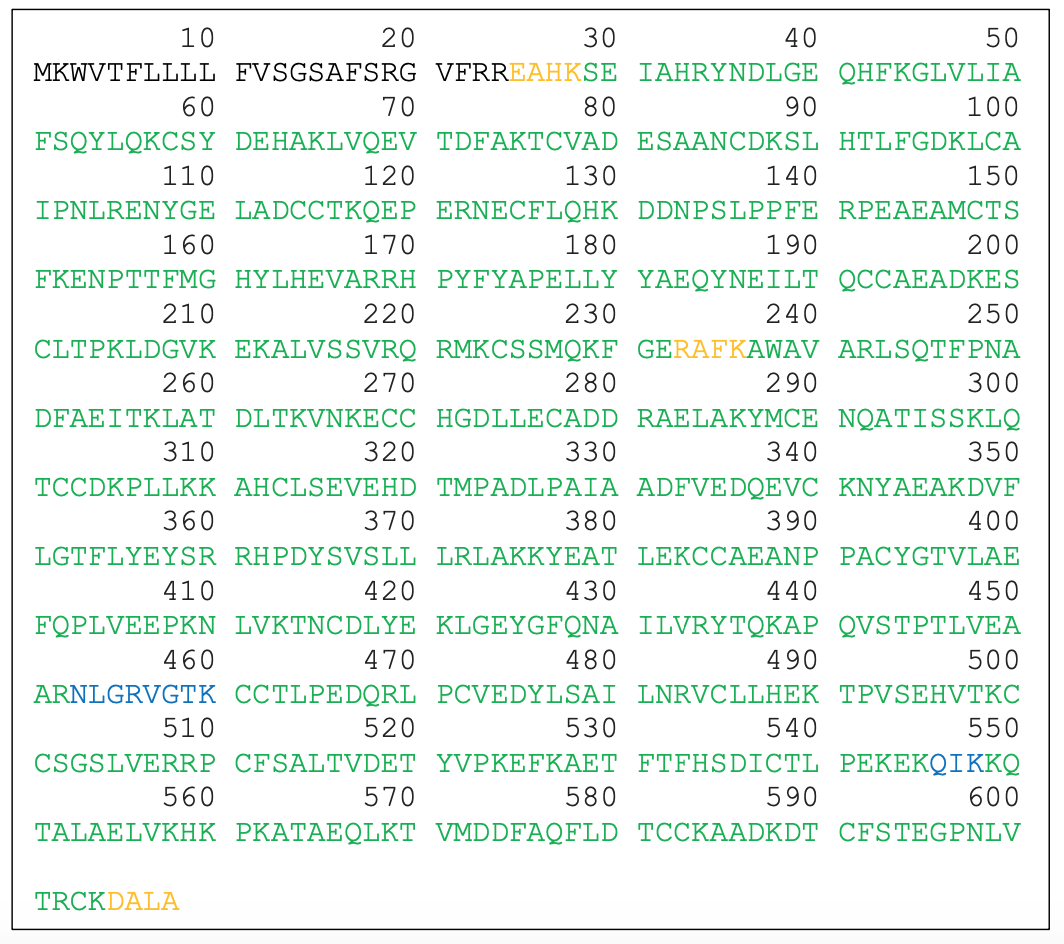

**Supplementary Figure 2.** FASTA amino acid sequence of mouse serum albumin (MSA, UniProt no. Q546G4) colored coded to the sequence coverage of MSA from digestion with trypsin or Lys-C and Glu-C determined by Maxquant. Green = covered by both digestions, yellow = only covered by trypsin digestion, blue = only covered by Lys- C/Glu-C digestion.

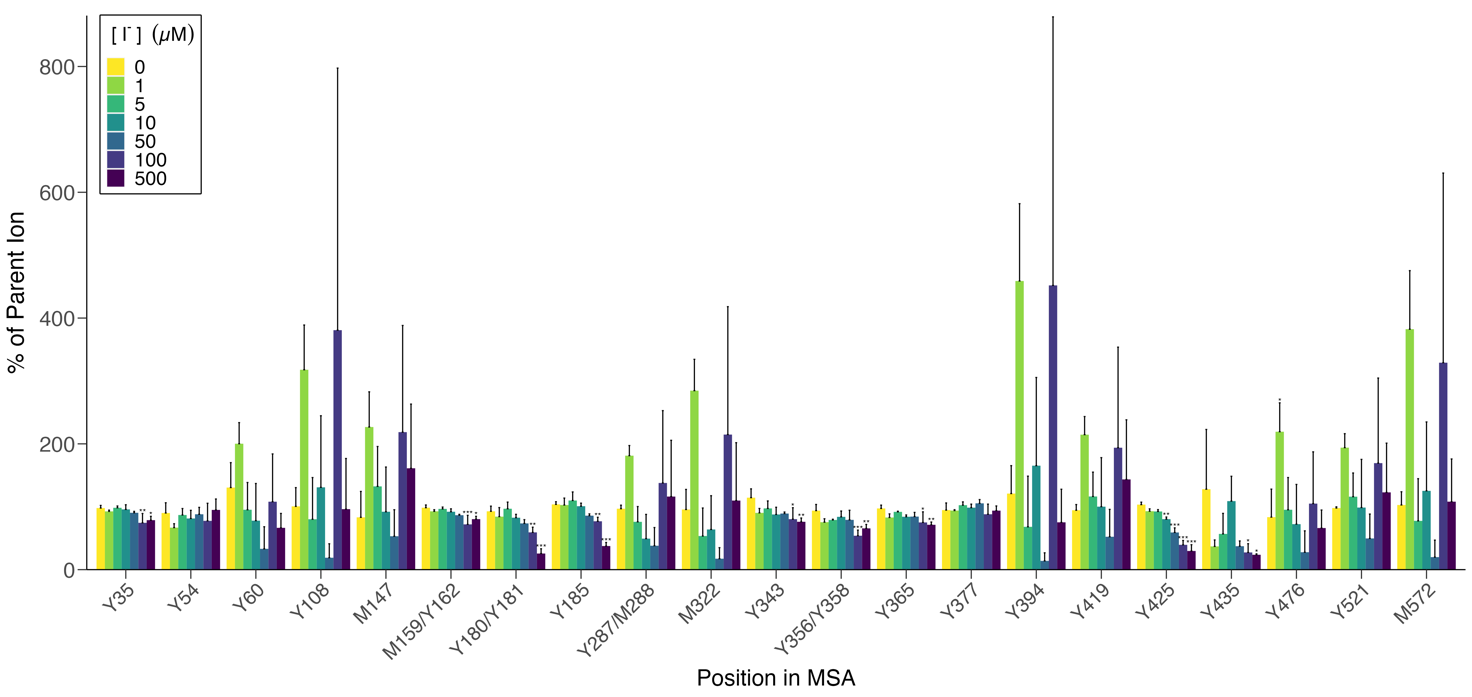

**Supplementary Figure 3**. Mouse serum albumin (5 µM) was treated with LPO (1.5 µM) and H_2_O_2_ (10x excess, 50 µM) in the absence or presence of increasing I⁻ concentrations (1 - 500 µM) for 2 h at 37 °C before reduction/alkylation, digestion and LC-MS/MS peptide mapping. Loss of parent (unmodified) peptide for all Tyr (Y) and Met (M) containing peptides that were identified as being modified. Some of the peptides contained two residues which could contribute to parent ion loss.

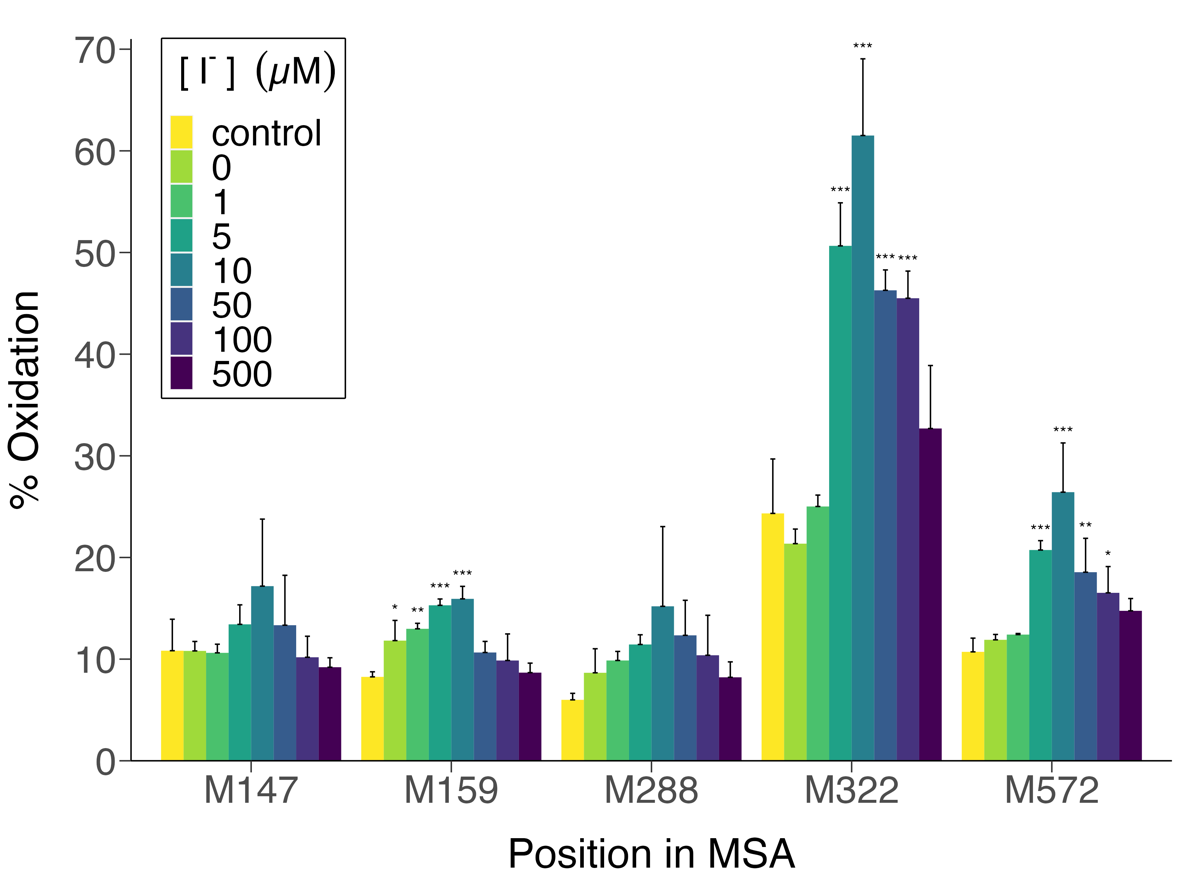

**Supplementary Figure 4.** Control samples of MSA contain significant amounts of oxidized Met (5 - 25% occupancy) but the extent of oxidation at M159, M322, M572 increased significantly on treatment with LPO/H₂O₂/I⁻ systems containing increasing levels of I⁻, though to variable extents. For reaction conditions see Supplementary Figure 3.

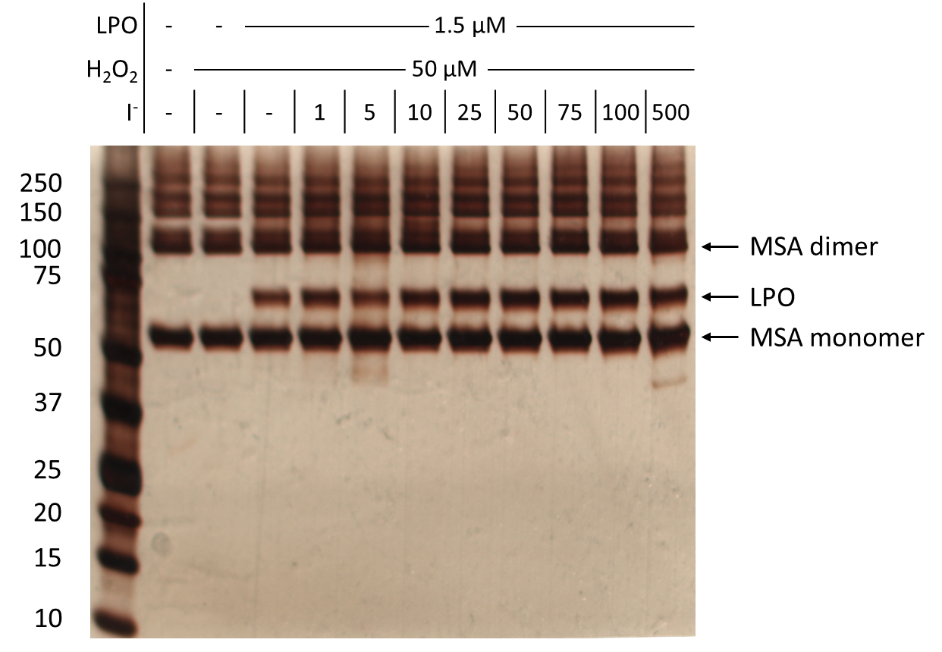

**Supplementary Figure 5.** Treatment of mouse serum albumin with LPO/H_2_O_2_/I⁻ system does not induce major structural changes as detected by SDS-PAGE with silver staining. Mouse serum albumin (5 µM) was treated with LPO (1.5 µM), H_2_O_2_ (10-fold molar excess over albumin, 50 µM), and I⁻ (1 - 500 µM) for 2 h at 37 °C in chelexed phosphate buffer, pH 5.8. Protein bands were detected by silver staining following SDS-PAGE separation under non-reducing conditions. The molecular masses of the markers present in the first lane are indicated at the left-hand side of the figure. The position of the bands assigned to MSA monomer, dimer and LPO are indicated on the right-hand side of the panel. The material above the MSA dimer band corresponds to higher aggregates present in the commercial samples of MSA with these detected in all samples.

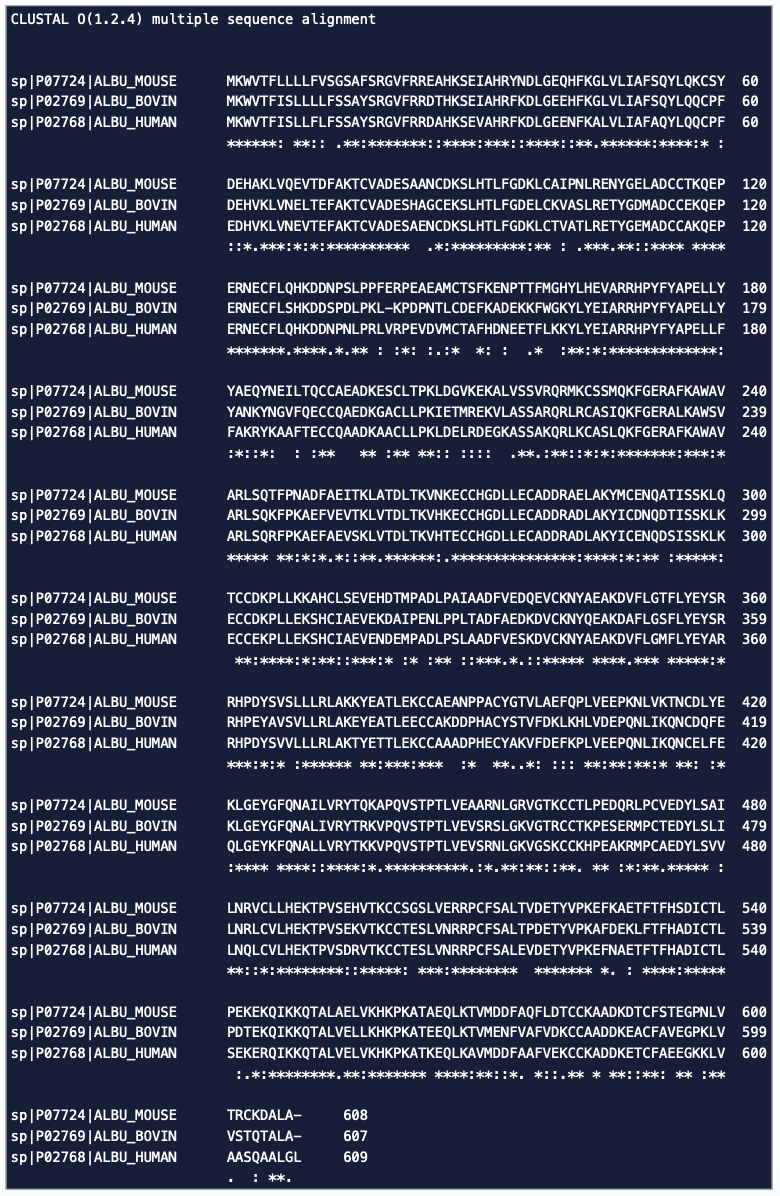

**Supplementary Figure 6.** Protein sequence alignment between human serum albumin (HSA, UniProt no. P02768), bovine serum albumin (BSA, UniProt no. P02769) and mouse serum albumin (MSA; UniProt no. P07724). * (asterisk) indicates identical amino acids, : (colon) indicates residues with highly similar properties, . (period) indicates residues with weakly similar properties. The alignment was done using the Clustal Omega program in UniProt.

MGNAPQPSHISKYILRWRPKNSVGRWKEATIPGHLNSYTIKGLKPGVVYEGQLISIQQYGHQEVTRFDFTTTSTSTPHHHHHH

**Supplementary Figure 6.** Translated sequence of the recombinant anastellin construct used in this study.

| **Condition** | **% Oxidation** | **SD** | **Significance** |
| --- | --- | --- | --- |
| control | 8.00 | 1.29 |  |
| 0 µM I^-^ | 10.1 | 0.516 |  |
| 1 µM I^-^ | 9.97 | 1.04 |  |
| 5 µM I^-^ | 11.4 | 1.33 | * |
| 10 µM I^-^ | 12.6 | 2.25 | ** |
| 50 µM I^-^ | 8.21 | 1.20 |  |
| 100 µM I^-^ | 6.49 | 1.32 |  |
| 500 µM I^-^ | 5.61 | 0.214 |  |

**Supplementary Table 1.** Quantification of the extent of modification (mono- and di-oxidation) of MSA at W238 on treatment with LPO/H₂O₂/I⁻ containing increasing levels of I⁻. MSA was treated with LPO (1.5 µM) and H_2_O_2_ (50 µM) in the absence and presence of increasing I⁻ concentrations (0 - 500 µM) for 2 h at 37 °C before trypsin digestion, reduction/alkylation, and analysis by LC-MS. Control = MSA alone. Trp oxidation was identified as a + 32 mass shift. Data are presented as mean and SD from 3 independent experiments. Statistical significance was determined by one-way ANOVA with a post-hoc Dunnett’s multiple comparison test. * p < 0.05, ** p < 0.01.

|  | **Anastellin** | **x** | **x** | **x** | **x** | **x** |
| --- | --- | --- | --- | --- | --- | --- |
|  | **MPO + H_2_O_2_ + Cl⁻** | **-** | **x** | **x** | **x** | **x** |
|  | **MPO + H_2_O_2_ + Cl⁻ + I⁻** | **-** | **-** | **10 μM** | **100 μM** | **1000 μM** |
| Peptide |  |  |  |  |  |  |
| GNAPQPSHISK |  | 100,0 | 99,4 | 97,7 | 89,5 | 75,0 |
| GNAPQPSH[+125.9]ISK |  | 0,0 | 0,4 | 2,2 | 9,2 | 20,7 |
| GNAPQPSH[+251.8]ISK |  | 0,0 | 0,2 | 0,1 | 1,3 | 4,3 |
| YILR |  | 99,9 | 93,3 | 63,7 | 36,6 | 17,9 |
| Y[+34]ILR |  | 0,0 | 6,3 | 0,5 | 0,0 | 0,0 |
| Y[+67.9]ILR |  | 0,1 | 0,4 | 0,0 | 0,1 | 0,0 |
| Y[+125.9]ILR |  | 0,0 | 0,0 | 29,4 | 50,2 | 56,0 |
| Y[+251.8]ILR |  | 0,0 | 0,0 | 6,4 | 13,1 | 26,0 |
| ATIPGHLNSYTIK |  | 100,0 | 54,7 | 43,5 | 21,3 | 17,0 |
| ATIPGHLNSY[+34]TIK |  | 0,0 | 24,5 | 4,0 | 0,0 | 0,0 |
| ATIPGHLNSY[+67.9]TIK |  | 0,0 | 20,7 | 0,0 | 0,0 | 0,1 |
| ATIPGHLNSY[+125.9]TIK |  | 0,0 | 0,0 | 31,1 | 28,2 | 28,2 |
| ATIPGHLNSY[+251.8]TIK |  | 0,0 | 0,0 | 21,4 | 50,5 | 54,8 |
| GLKPGVVYE |  | 100,0 | 78,8 | 91,6 | 90,4 | 80,8 |
| GLKPGVVY[+34]E |  | 0,0 | 18,7 | 2,4 | 0,0 | 0,0 |
| GLKPGVVY[+67.9]E |  | 0,0 | 2,4 | 0,1 | 0,0 | 0,0 |
| GLKPGVVY[+125.9]E |  | 0,0 | 0,0 | 5,8 | 9,1 | 17,1 |
| GLKPGVVY[+251.8]E |  | 0,0 | 0,0 | 0,1 | 0,5 | 2,0 |
| GQLISIQQYGHQE |  | 100,0 | 89,1 | 40,5 | 18,3 | 16,6 |
| GQLISIQQY[+34]GHQE |  | 0,0 | 9,4 | 0,4 | 0,0 | 0,0 |
| GQLISIQQY[+67.9]GHQE |  | 0,0 | 1,5 | 0,0 | 0,0 | 0,0 |
| GQLISIQQY[+125.9]GHQE |  | 0,0 | 0,0 | 28,3 | 30,8 | 32,6 |
| GQLISIQQY[+251.8]GHQE |  | 0,0 | 0,0 | 30,8 | 50,9 | 50,7 |

**Supplementary Table 2** Mono- and dichlorinated and iodinated peptides detected in anastellin exposed to MPO in the presence and absence of iodide followed by proteolytic digestion with trypsin and Glu-C. Anastellin (5 µM) was treated with MPO (0.1 µM), H_2_O_2_ (50 µM, 10-fold molar excess over protein), Cl⁻ (100 mM), and I⁻ (0 - 1000 µM) for 2 h at 37 °C. The relative occupancy is calculated by dividing the area-under-the-curve for a particular species with the total area-under-the-curve for all the detected species of the corresponding peptide. Data are presented as a mean from 3 independent experiments.
